# Stochastic gene expression is optimized to drive developmental self-organization

**DOI:** 10.1101/546911

**Authors:** Ritika Giri, Dimitrios K. Papadopoulos, Diana M. Posadas, Hemanth K. Potluri, Pavel Tomancak, Madhav Mani, Richard W. Carthew

**Affiliations:** Department of Molecular Biosciences, Northwestern University, Evanston, IL 60208, USA; NSF-Simons Center for Quantitative Biology, Northwestern University, Evanston, IL 60208, USA; Max Planck Institute of Molecular Cell Biology and Genetics, Dresden, Germany; Department of Engineering Sciences and Applied Mathematics, Northwestern University, Evanston, IL 60208, USA; Department of Biochemistry and Molecular Genetics, Northwestern University, Chicago, IL 60611, USA

**Author notes:** MRC Human Genetics Unit, Institute of Genetics and Molecular Medicine, University of Edinburgh, UK.

## Abstract

Self-organization of cells into tissue patterns is a design principle in developmental biology to create order from disorder. However, order must emerge from biochemical processes within and between cells that are stochastic. Here, we measure noise in expression of the *Drosophila senseless* gene, a key determinant of sensory cell fate. We show that translation and transcription of *senseless* produce distinct signatures of protein noise. Repression of *senseless* by a microRNA uniformly decreases both protein abundance and noise in cells, but does so without affecting the fidelity of self-organization. In contrast, the genomic location of *senseless* affects protein noise without affecting protein abundance. When noise is greater, tissue patterning is significantly disordered. This suggests that gene expression stochasticity, independent of expression level, is a critical feature that must be constrained during development to allow cells to efficiently and accurately self-organize.

---

Stochasticity is a fundamental feature of all molecular interactions. During gene expression, stochasticity leads to fluctuations in the number of protein molecules in a cell^1^ These fluctuations are perhaps especially relevant during developmental transitions, when cells adopt divergent fates based on the protein abundance of a fate determinant. While probabilistic fate adoption has been observed in rare instances^2–4^, generally, fate transitions are thought to be virtually deterministic due to cell lineage and proximity to inductive signals. Therefore, it is unclear how extensively protein fluctuations impinge upon the vast majority of developmental decisions. Given that the emergence of order from disorder is a hallmark of development, how are highly-ordered patterns rendered immune to the underlying stochasticity of gene expression? We explore this problem by focusing on the patterning of sensory bristles in *Drosophila*.

Sensory organs are often arranged in highly-ordered assemblies, as a means to predictably map environmental stimuli to neural circuitry^5^. Paradoxically, *Drosophila* sensory bristle development requires expression heterogeneity between progenitor cells to initiate the self-organizing process of pattern formation^6,7^ We have focused on sensory bristles located at the anterior margin of the wing, where diffusing Wingless (Wg) molecules induce the expression of *senseless* (*sens*) in stripes of cells. This imbues these cells with a bistable fate potential (Fig. 1a)^8^. Cells then either upregulate *sens* expression and adopt a sensory organ (S) fate or downregulate *sens* and adopt an epidermal (E) fate^9,10^ Cells in each stripe compete with one another to adopt an S fate, which is driven by lateral inhibition of *sens* expression via Notch signaling^7^ Sustained expression of the Sens transcription factor is sufficient to drive a cell towards an S fate, after which the cell develops into an adult bristle (Fig. 1b)^9^. This key role of Sens^11^ makes it a logical candidate to study how quantitative changes in gene expression noise impact the final ordered outcome.

**Figure 1.**
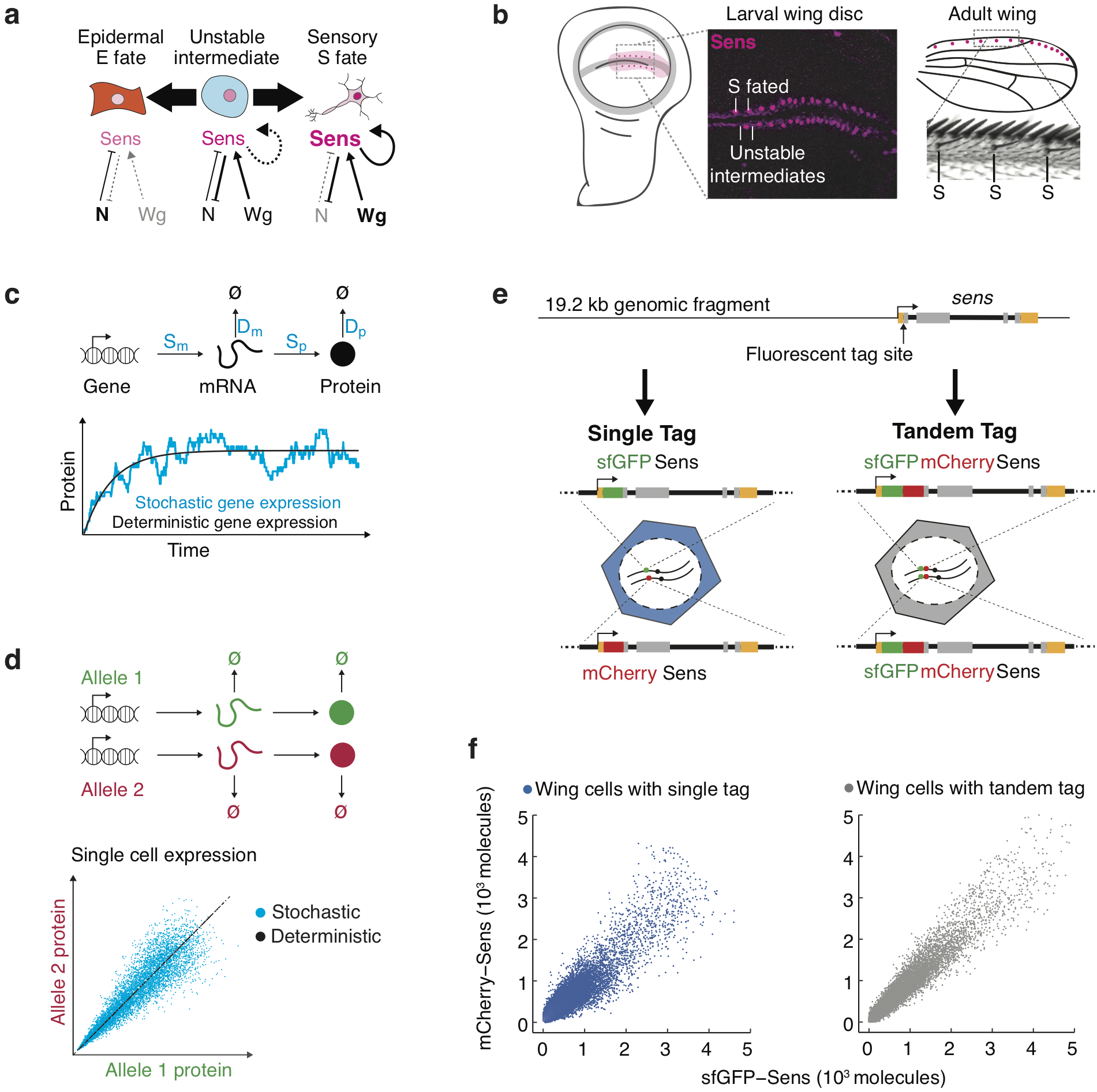
Measuring *in vivo* gene expression stochasticity during sensory organ fate selection. **a,** Sensory organ (S) fate selection is a bistable system driven by pro-neural Wingless (Wg) and anti-neural Notch (N) signals to regulate *sens* expression. When cells are in an unstable state of intermediate expression, they switch to either low stable expression (E fate) or high stable expression (S fate). **b,** Sens expression marks the resolving proneural field along the *Drosophila* wing margin. The stripes of unstable intermediate cells refine into a robust periodic pattern of S cells, which can be seen emerging in the micrograph, and E cells, which will emerge from the unstable intermediates interspersed between S cells. This generates the bristle pattern along the adult wing margin. **c,** Gene expression output is inherently variable due to stochastic synthesis and decay of mRNA and protein molecules. Therefore, single cell protein counts fluctuate stochastically from the deterministic expectation. The magnitude of these fluctuations is determined by the rate constants of individual steps (in blue). **d,** Stochasticity can be measured by tagging the two alleles of a gene with distinct fluorescent proteins and measuring fluorescence correlation in individual cells. Perfect correlation would indicate no stochastic effects i.e. deterministic behavior. **e,** A genomic fragment containing *sens* was N-terminally tagged with either single sfGFP or mCherry tags; or with a tandem sfGFP-mCherry fluorescent tag. These were used to rescue *sens* null animals by site-specific insertion into genomic location 22A3. **f,** Single-cell mCherry and sfGFP protein was measured in singly tagged *sfGFP-sens/mCherry-sens* wing cells. Tandem tagged *sfGFP-mCherry-sens* wing cells serve as a technical control for non-biological sources of stochasticity.

## Measuring stochasticity in *sens* expression

Protein fluctuations can be observed by counting molecules in single cells over time (Fig. 1c)^12^. Alternatively, noise can be estimated by tagging the two alleles of a gene with distinct fluorescent proteins and measuring fluorescence correlation in individual cells (Fig. 1d)^1,13^. We modified a 19.2 kb fragment of the *Drosophila* genome containing *sens* by singly inserting either superfolder GFP (sfGFP) or mCherry into the amino terminus of the *sens* ORF^14^. These transgenes were precisely landed into the 22A3 locus on the second chromosome, a standard landing site for transgenes (Fig. 1e). Endogenous *sens* was eliminated using protein-null alleles^9,10,15^. The transgenes completely rescued all detectable *sens* mutant phenotypes and exhibited normal expression (Extended Data Fig. 1). We then mated singly-tagged *sfGFP-sens* with *mCherry-sens* animals to generate heterozygous progeny. Wing imaginal discs of these offspring were fixed and imaged by confocal microscopy (Extended Data Fig. 2). Cells were computationally segmented in order to measure intensity of sfGFP and mCherry fluorescence within each cell (Extended Data Fig. 3). Fluorescence values were then converted into absolute numbers of Sens protein molecules. This was made possible by using Fluorescence Correlation Spectroscopy (FCS) to measure the concentrations of sfGFP-Sens and mCherry-Sens protein in live wing discs and deriving a conversion factor from these measurements (Extended Data Fig. 4).

Wing disc cells displayed a wide range of Sens protein expression (Fig. 1f). This was expected since *sens* transcription is regulated by Wg and Notch signaling^8,9^ Although both sfGFP and mCherry tagged alleles contributed equally to total Sens output on average (Extended Data Fig. 4a), individual cells showed significant differences between sfGFP and mCherry fluorescence (Fig.1f). This was due to two independent sources of noise: 1) stochastic gene expression, 2) stochastic processes linked to imaging and analysis. The latter arises from differential fluorescent maturation kinetics, probabilistic photon emission and detection, as well as image analysis errors. To estimate this technical noise, we constructed a third transgene containing both sfGFP and mCherry fused in tandem to the *sens* ORF (Fig. 1e). *sfGFP-mCherry-sens* was inserted at locus 22A3, and fluorescence was measured in disc cells from such animals. Since sfGFP and mCherry molecule numbers should be perfectly correlated *in vivo* when expressed as a tandem-tagged protein, we attributed any decrease in fluorescence correlation to technical noise (Fig. 1f).

Previous studies using dissociated cells have shown that protein output from gene expression is Gamma-distributed^1,16^, such that protein noise (expressed as the Fano factor, i.e. variance divided by mean) remains constant as protein output varies. We calculated the Fano factor as a function of Sens protein output in cells expressing either singly-tagged Sens or tandem-tagged Sens (Fig. 2a). To estimate the Fano factor due to stochasticity of Sens expression, we subtracted out the technical contribution as measured in tandem-tagged cells. Contrary to expectation, the Fano factor for Sens expression displayed a complex relationship to protein output, with a peak in cells containing < 300 molecules (Fig. 2b).

**Figure 2.**
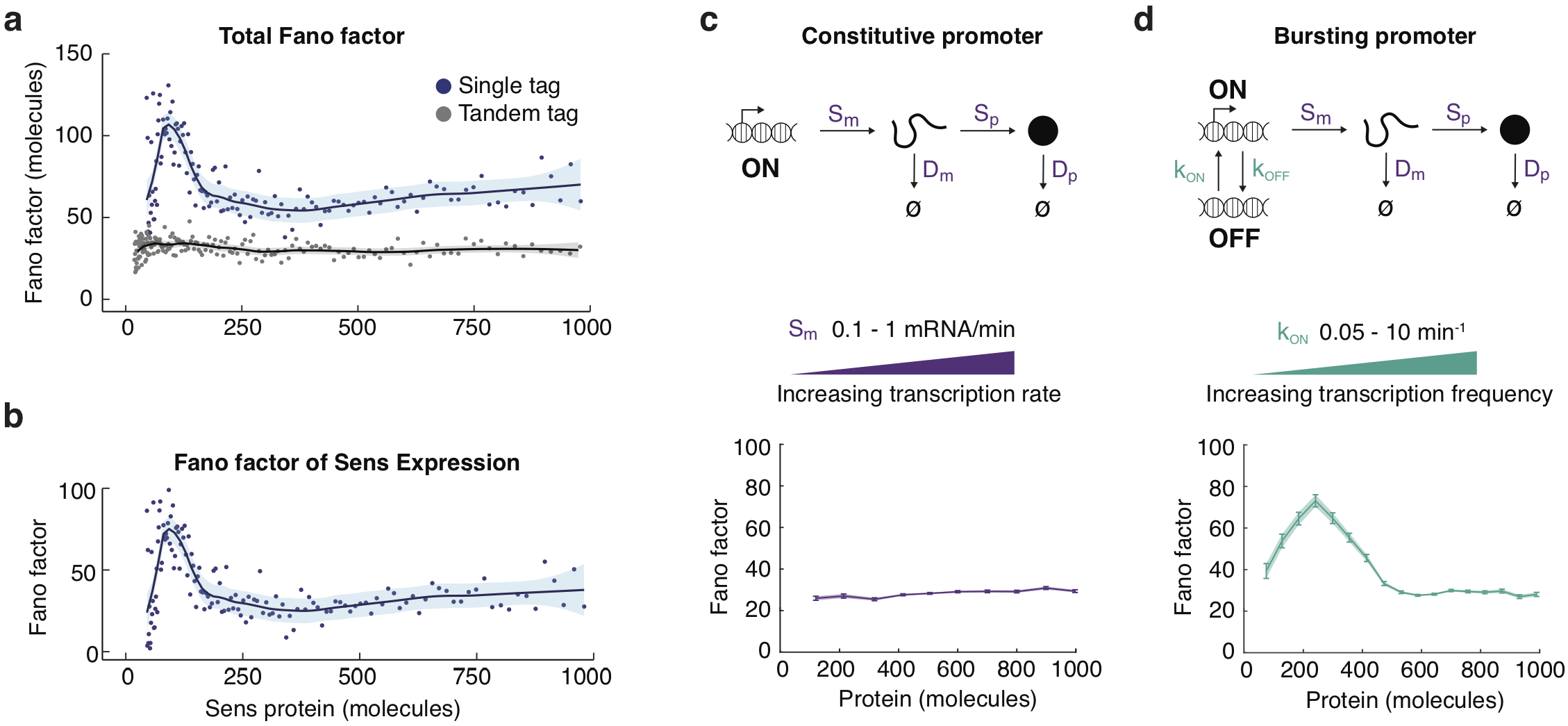
Sens protein expression displays signatures of transcriptional bursting. **a,** The Sens Fano factor as measured in bins of wing disc cells expressing either tandem-tagged Sens or the singly-tagged allelic pairs of Sens. **b,** The Fano factor of Sens expression was calculated by subtracting out the Fano factor from tandem-tagged cells. For **a** and **b,** shading demarcates 95% confidence intervals. **c,** Simulations of a gene expression model with a constitutively-active promoter produces a protein Fano factor that is invariant with protein output, like a Gamma process. Transcription rate *S_m_* is being varied from 0.1 to 1 mRNA/min to generate graded Sens expression. **d,** Simulations of a gene expression model with a bursting promoter with distinct on and off states, and independent rate constants for state conversion. If the rate constant *k_on_* is varied as shown, the simulated protein Fano factor exhibits a biphasic profile as a function of protein output. Above a threshold of protein output, the Fano factor becomes invariant. This complex behavior is observed experimentally in **b.** For **c** and **d,** error bars represent 95% confidence intervals.

## Numerical modeling of *sens* gene expression

To understand the origins of this profile, we created a mathematical model of gene expression using rate parameters that we measured for *sens* in the wing (Supplementary Table 2). Each reaction in the model was treated as a probabilistic event, reflecting the stochastic nature of these processes^17,18^. Thus, simulated protein levels displayed fluctuations. To mimic the experimental set-up, two independent alleles were simulated for each virtual cell. *In silico*, the results resembled the *sens* allelic output that we observed *in vivo* (Extended Data Fig. 5).

We first considered a model in which the promoter was always active. The simulated Fano factor was constant irrespective of protein number, consistent with noise being caused by random birth-death events (Fig. 2c)^17,18^. This resembled the *in vivo* profile seen in cells containing more than 300 molecules of Sens (Fig. 2b). Since there was a weak rise in the experimental Fano factor, we hypothesize that one of the post-transcriptional rate constants slowly varies as a function of protein output (Extended Data Fig. 5d). Notably, the model did not recapitulate the observed Fano peak at lower Sens levels. Thus, Sens fluctuations were not exclusively due to the random birth and death of mRNA and protein molecules.

Therefore, we considered an alternative model (Fig. 2d). The promoter was allowed to switch between active and inactive states such that it transcribed mRNA molecules in bursts. Bursty transcription is a common feature of gene expression in many organisms^19–21^. Varying the three transcription parameters in our model (*k_on_, k_off_*, and *S_m_*) allowed us to independently tune the frequency (*k_on_*^−1^ + *k_off_*^−1^)^−1^ and size (*S_m_/k_off_*) of these virtual bursts. When we systematically varied the gene activation parameter *k_on_* and calculated the resulting Fano factor, the *in-silico* profile closely resembled the *in vivo* profile (Fig. 2d). In contrast, varying inactivation rate *k_off_* (Extended Data Fig. 5f) or transcription rate *S_m_* (Fig. 2c) did not yield a Fano peak at lower Sens levels. We surmise from these results that transcription of *sens* at the wing margin is primarily regulated by modulating promoter burst frequency via *k_on_*. Indeed, burst frequency modulation has been observed for multiple developmental genes^22^ and might be a conserved mechanism to reduce stochastic noise.

These simulations allow us to conceptually frame Sens protein noise as coming from two distinct sources 1) transcriptional bursting kinetics, and 2) RNA/protein birth-death processes. When transcription bursts are infrequent, cells experience large fluctuations in mRNA and protein numbers, which generates a peak in the protein Fano factor. As promoter activation events become more frequent, they approximate a continuous-rate process such that RNA/protein birth-death processes dominate the protein noise, and the Fano factor becomes more constant.

## miR-9a modestly suppresses expression noise

The magnitude of the Fano factor should be proportional to the average number of protein molecules translated from one mRNA molecule in its lifetime, if protein noise is caused by RNA/protein birth-death processes^17,18,23,24^. Therefore, we experimentally altered this translation burst size for Sens. We did so by eliminating the post-transcriptional repression of Sens by the microRNA miR-9a (Fig. 3a)^15,25^. Loss of miR-9a repression increased Sens protein numbers per cell by 1.8-fold (Fig. 3b, Extended Data Fig. 6a). We then compared the Fano factor in cells with or without miR-9a regulation. Loss of miR-9a regulation increased the Fano factor across the entire range of *sens* expression (Fig. 3c). We compared this effect to model simulations in which either the translation rate (*S_p_*) or mRNA decay rate (*D_m_*) parameter was altered by 1.8-fold. The model-predicted increase in the Fano factor was comparable to the observed elevation when miR-9a regulation was missing (Fig. 3d).

**Figure 3.**
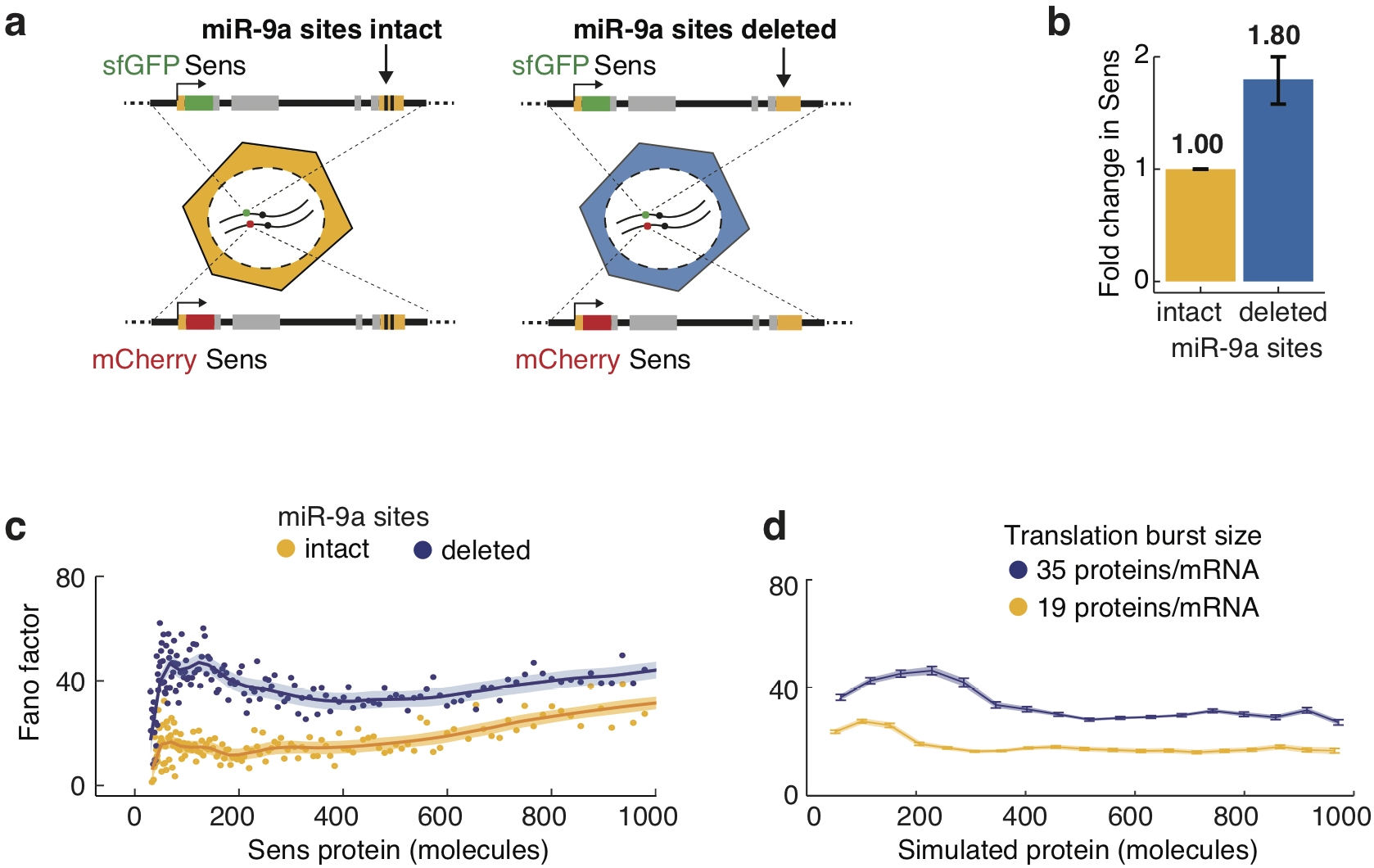
MicroRNA regulation decreases Sens protein output and noise. **a,** The *sens* transgenes were modified to delete the two miR-9a binding sites in the 3’UTR. Sens output was measured in singly tagged allele pairs with and without miR-9a sites. **b,** Sens protein output increased 1.80 ± 0.21 fold with loss of miR-9a repression. Error bars are standard error of the mean. **c,** Loss of miR-9a regulation leads to an increase in Fano factor across the entire range of protein output. Shaded regions are 95% confidence intervals. **d,** Model simulations with a 1.8 fold increase in translation burst size (defined as *S_p_/D_m_*) reproduces the experimental Fano profile. Error bars are 95% confidence intervals.events become more frequent, they approximate a continuous-rate process such that RNA/protein birth-death processes dominate the protein noise, and the Fano factor becomes more constant.

## Inter-allelic gene transvection dramatically increases Sens protein noise

Our model had suggested that promoter switching is the dominant source of protein noise in cells with fewer than 300 Sens molecules. To test this hypothesis, we landed the *sens* transgenes in a different location of the genome (Fig. 4a). We reasoned that a different genomic neighborhood would possibly alter promoter bursting dynamics. We chose 57F5 to land *sens*, since both 22A3 and 57F5 are widely used landing sites for *Drosophila* transgenes^14^. The *sens* transgenes inserted at 57F5 were highly comparable to the 22A3 site in their ability to rescue null *sens* mutations, as well as express Sens protein in the correct pattern and abundance at the wing margin (Extended Data Fig. 2).

**Figure 4.**
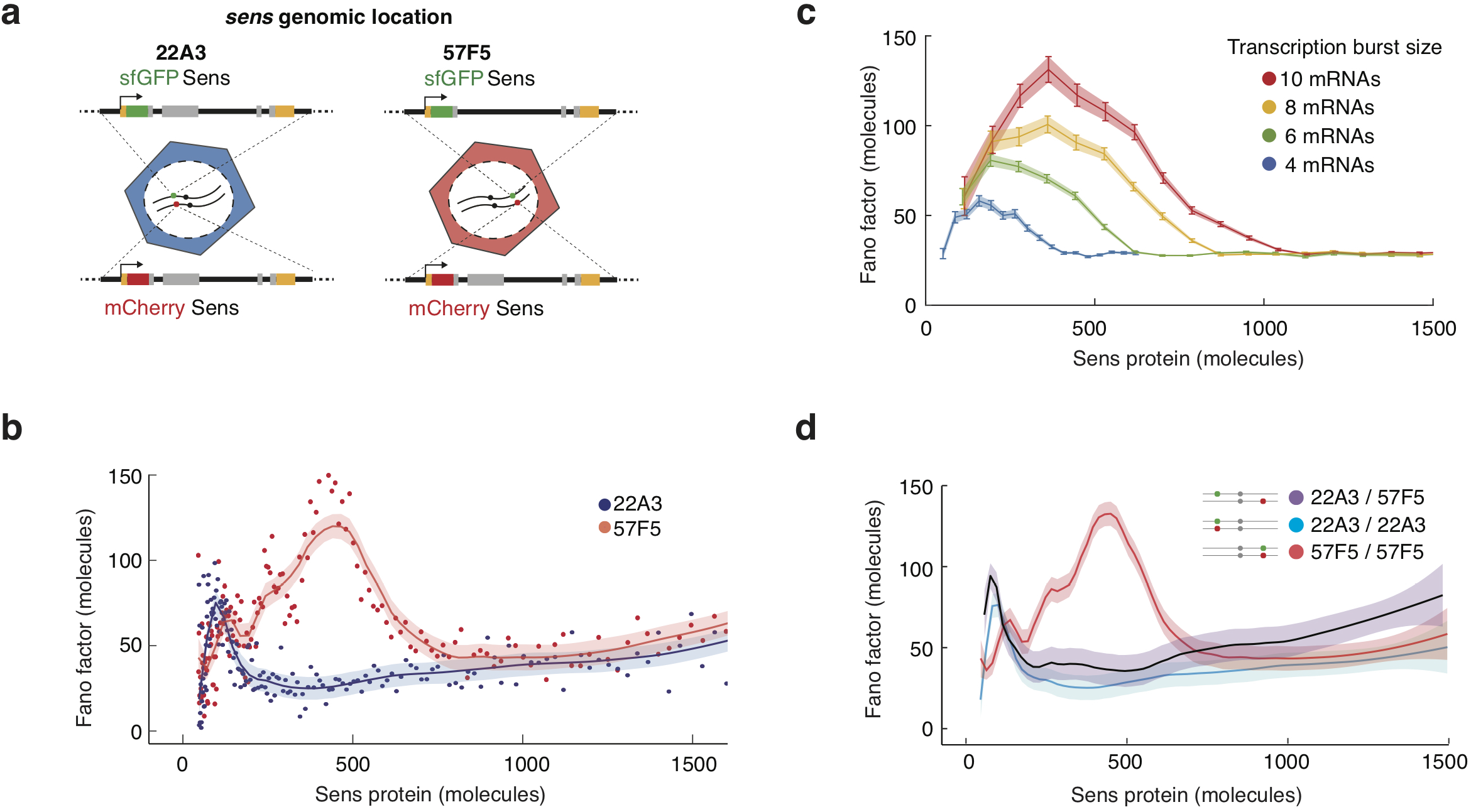
Sens protein noise is dependent on genome location of the *sens* gene. **a,** The *sens* transgenes were inserted into one of two locations on chromosome II - 22A3 or 57F5. **b,** The noise profiles from cells expressing *sens* at 57F5 or 22A3. Cells with more than 800 molecules have identical Fano values at both genomic positions. The Fano peaks at lower Sens levels are very different between the genomic locations. Shaded regions are 95% confidence intervals. **c,** Model simulations in which transcription burst size (defined as *S_m_/k_off_*) is set to different values as shown. The Fano peak amplitude and position change as burst size varies, but all relax to a constant basal level. These trends are highly similar to those observed in **b.** Error bars are 95% confidence intervals. **d,** The noise profiles from cells expressing sens as paired alleles (22A3/22A3, 57F5/57F5) or unpaired alleles (22A3/57F5). Line averages are shown and shaded regions are 95% confidence intervals.

Cells expressing Sens from 57F5 and containing more than 800 Sens molecules had a Fano factor that was identical to their 22A3 counterparts (Fig. 4b). However, the profile was very different in cells with fewer than 800 molecules; the Fano factor peak was larger and greatly expanded. It was remarkable how different the profiles appeared, and we explored potential mechanistic causes of these differences. When varying the transcriptional parameters in the model we found that even a modest increase in burst size could recapitulate the effect of changing gene location from 22A3 to 57F5 on noise (Fig. 4c).

We looked for local properties of the genome that might explain the difference between 22A3 and 57F5. Metazoan genomes are segregated into Topologically Associated Domains (TADs). TADs are conserved 3D compartments of self-interacting chromatin whose boundaries are demarcated by insulator protein binding. TADs often differ in chromatin condensation and accessibility to transcription factors^26^. The landing site at 22A3 is in the middle of a large, inaccessible, gene-sparse TAD (Extended Data Fig. 7)^27^. In contrast, the landing site at 57F5 is in a small, accessible, gene-dense TAD. Moreover, the 22A3 site is 40 kb from bound insulators while the 57F5 site is located less than 1 kb from a TAD boundary. Proximity to insulator elements is associated with transcriptional interactions between paired alleles of *Drosophila* genes^22,28,29^. To test whether altered transcriptional kinetics of *sens* at 57F5 were allele-intrinsic or due to inter-allelic interactions, we placed a 57F5 allele *in trans* to a 22A3 allele, generating unpaired alleles. The Fano factor profile of 22A3/57F5 cells was strikingly similar to that of 22A3/22A3 cells (Fig. 4d). In contrast, the model predicted that if the alleles behaved intrinsically, then the Fano factor of 22A3/57F5 cells would have been much greater than that of 22A3/22A3 cells (Extended Data Fig. 5g). This suggests that alleles paired at 22A3 frequently fire independently of one another, but alleles paired at 57F5 exhibit inter-allelic interactions, also called transvection. Loss of homolog pairing in 22A3/57F5 cells presumably reverts the Fano factor profile to mimic the non-interacting 22A3 pair (Fig. 4d).

## Sens noise can disrupt patterning order of adult bristles

Although we observed dramatic changes in Sens noise when we altered genomic location, heightened fluctuations were limited to cells with fewer than 800 Sens molecules, far lower than the level of Sens expression in S-fated cells. Therefore, it is possible that these fluctuations do not impact fate transitions and bristle patterning. Remarkably, this is not the case. Instead, we find that increased Sens stochasticity in this regime results in increased pattern disorder in the adult form.

We had measured Sens noise in cells undergoing fate decisions to make chemosensory bristles. Chemosensory bristles are periodically positioned in a row near the adult wing margin, such that approximately every fifth cell is a bristle (Fig. 5a)^30^. Mechanosensory bristles form in a continuous row at the outermost margin of the adult wing and are selected 8-10 hours after the chemosensory cells are selected^30^. We reasoned that if Sens numbers fluctuate in sensory progenitor cells near the bistable switching threshold, it might propel erroneous escape from lateral inhibition to generate ectopic sensory organs. Thus, mechanosensory bristles positioned incorrectly in the chemosensory row might be derived from proneural cells that escaped inhibition during chemosensory specification (Fig. 5a). Indeed, when we compared ectopic bristles in 57F5 versus 22A3 adults, the frequency increased ten-fold from 3.2% to 29.1% in 57F5 adults (Fig. 5b, Supplementary Table 3). Ectopic chemosensory bristles were also observed (Figs. 5a, Extended Data Fig. 8). The difference in errors was not attributable to genetic background in the different lines since parental stocks had identical ectopic bristle frequencies (Fig. 5b, Extended Data Fig. 8). Nor was it due to higher Sens protein levels in cells from 57F5 animals since there was no dramatic difference in the Sens levels between 57F5 and 22A3 cells (Extended Data Fig. 2). Moreover, loss of miR-9a regulation increased Sens levels (Fig. 3b) but did not increase ectopic bristle frequency (Fig. 5c, Supplementary Table 3). Finally, we ruled out the possibility that insertion near neurogenic genes was responsible for enhanced ectopic bristles from 57F5. First, none of the genes residing in the 57F5 TAD are annotated as neurogenic (Supplementary Table 4)^15^. Second, adults with *sens* at 22A3/57F5 had an ectopic frequency of 1.7%, not significantly different from 22A3/22A3 adults (Fig. 5b, Supplementary Table 3).

**Figure 5.**
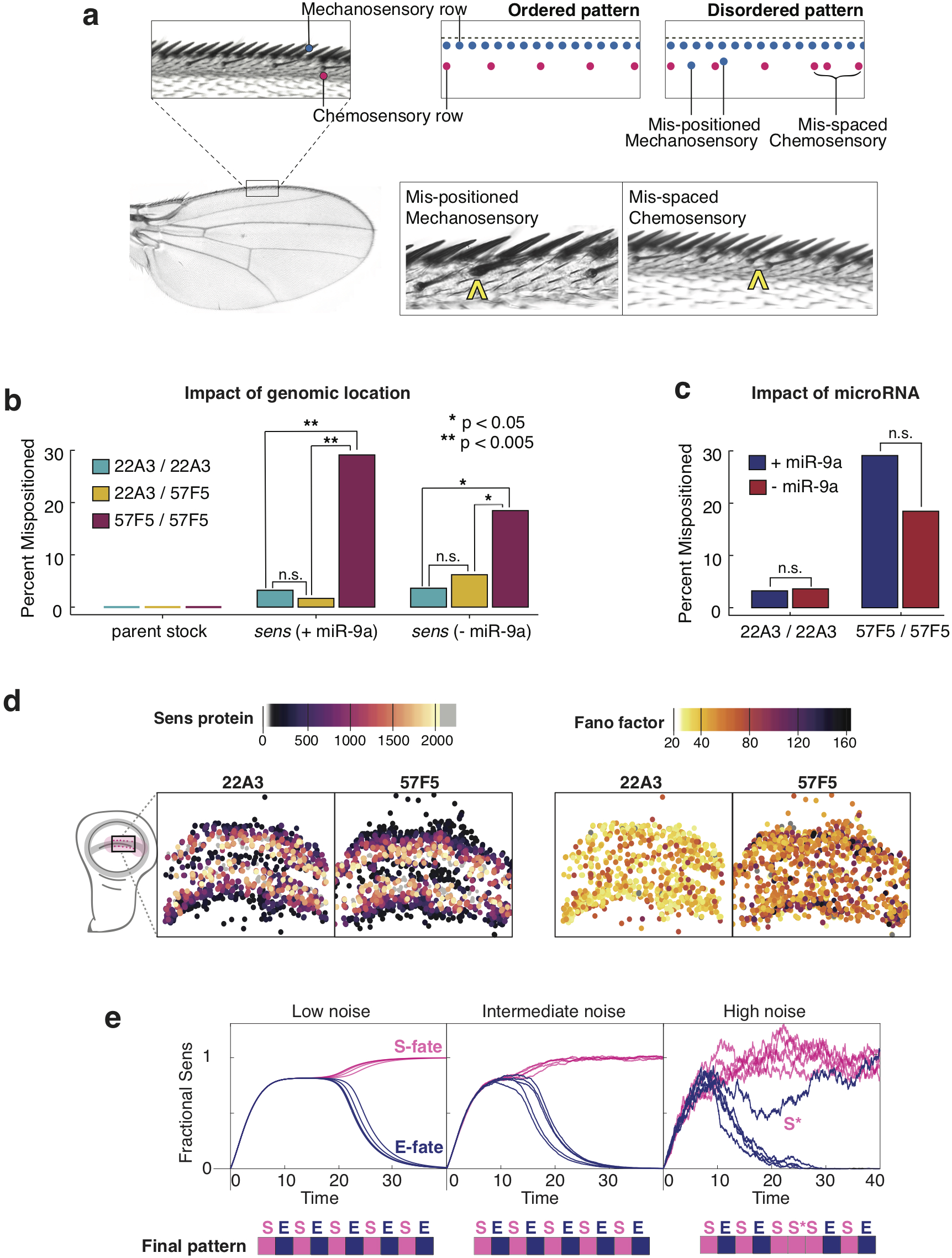
Ordered sensory patterning is disrupted by stochastic gene expression. **a,** The dorsal surface of the adult wing margin displays two ordered rows of sensory organs - an outer continuous row of thick mechanosensory bristles (cyan) and an inner periodic row of thin chemosensory bristles (magenta). Disordered patterns are observed when bristles are incorrectly positioned. Instances of ectopic (mis-positioned) mechanosensory bristles in the chemosensory row (center) and ectopic chemosensory bristles, which disrupt periodic spacing (right), were observed and counted. **b,** Pattern disorder is significantly higher when *sens* allele pairs are expressed from locus 57F5/57F5 relative to alleles at 22A3/22A3 or 22A3/57F5. This effect is seen regardless of whether miR-9a regulates *sens* or not. **c,** Increasing mean Sens levels 1.8-fold by removing miR-9a regulation does not lead to increased mispatterning events at either locus. **d,** The centroids of Sens-positive cells in 22A3/22A3 and 57F5/57F5 wing discs were mapped and color coded according to Sens expression level (left) and Fano noise level (right). Cells experiencing high Fano noise were distributed throughout the proneural zone of the 57F5 disc where S-fate determination occurs. **e,** Quantitative model of Sens protein dynamics during pattern formation. Sample simulations use a ID model adapted from Corson et al.,(2017)^7^ that generates periodic E and S cells as a final pattern. When all cells first express Sens, they enter an unstable state in which Sens levels are intermediate. Cells then bifurcate into alternating states with maximal or minimal Sens. The magnitude of noise in Sens expression affects the time taken to resolve the pattern and the accuracy of the pattern.

To understand why 57F5 cells with relatively low Sens numbers and high Sens noise disrupted patterning order, we mapped the location of these cells in the wing disc. Wg induces Sens in two broad stripes of cells, each stripe being 4-5 cell diameters wide (Fig. 5d)^8,31^. It is only cells near the center of a stripe that express higher levels of Sens^7,31^, and a few of these will switch to an S fate. This pattern was preserved whether *sens* was transcribed from 22A3 or 57F5 (Fig. 5d, Extended Data Figure 9a). However, the pattern of noise was remarkably different. For the 22A3 gene, cells with the greatest noise were at the edges of each stripe, distant from the central region from which S cells normally emerge (Fig. 5d, Extended Data Figure 9b). In contrast, for the 57F5 gene, cells with high noise were located throughout the stripe, including the central region. Thus, it is likely that a subset of cells encompassed in the Fano peak for the 57F5 gene were close to the bistable switch threshold and therefore susceptible to errors in fate determination due to the enhanced fluctuations in Sens.

Noise in *sens* expression appears to be a fine-tuned parameter. If noise is too low, then the final bristle pattern will be highly ordered, but cells will take more time to resolve their fates since noise initiates the self-organizing process of pattern formation (Fig. 5e). If noise is too high, then cells will rapidly resolve their fates, but the final pattern will be disordered (Fig. 5e). In this high noise scenario, a cell may experience a random fluctuation large enough to flip the cell into an inappropriate stable state during the resolution of pattern formation. Overall, it suggests that perhaps stochasticity in gene expression is an evolutionarily constrained parameter that allows rapid yet accurate cell fate resolution without requiring large numbers of fate-determining molecules^32–34^.

## Acknowledgments

Fly stocks from Hugo Bellen and the Bloomington Drosophila Stock Center are gratefully appreciated. Antibodies were gifts from Hugo Bellen and purchases from the Developmental Studies Hybridoma Bank. We thank Koen Venken for extensive help with BAC recombineering protocols and reagents. We thank Michael Stadler and Michael Eisen for generously providing HiC maps of the 22A3 and 57F5 loci from their studies. We thank Jessica Hornick and the Biological Imaging Facility for help with imaging, and Lionel Fiske for help implementing the fate selection model.

### Funding

Financial support was provided from the Northwestern Data Science Initiative (R.G.), Robert H. Lurie Comprehensive Cancer Center (R.G.), Pew Latin American Fellows Program (D.M.P.), Max Planck Society (D.K.P. and P.T.), Chancellors Fellowship of University of Edinburgh (D.K.P.), NIH (R35GM118144, R.W.C.), NSF (1764421, M.M and R.W.C.), and the Simons Foundation (597491, M.M. and R.W.C.). M.M. is a Simons Foundation Investigator.

### Author contributions

The experimental work was conceived by R.G. and R.W.C. D.M.P. and H.P. helped R.G. in the recombineering of *sens* transgenes. Fluorescence correlation spectroscopy was performed by D.K.P., under the supervision of P.T. All other experimental work was performed by R.G., under the supervision of R.W.C. Mathematical data analysis and modeling was conceived by R.G. and M.M. All data and model analysis was performed by R.G. under the supervision of M.M. The manuscript was written by R.G. and R.W.C. with input from all authors.

### Competing interests

Authors declare no competing interests.

### Data and materials availability

All stocks and molecular reagents will be made freely available upon request. Raw data of segmented cell fluorescence and position for all experiments as well as R and Matlab code used for analysis is available on request.

---

**Extended Data Figure 1.**
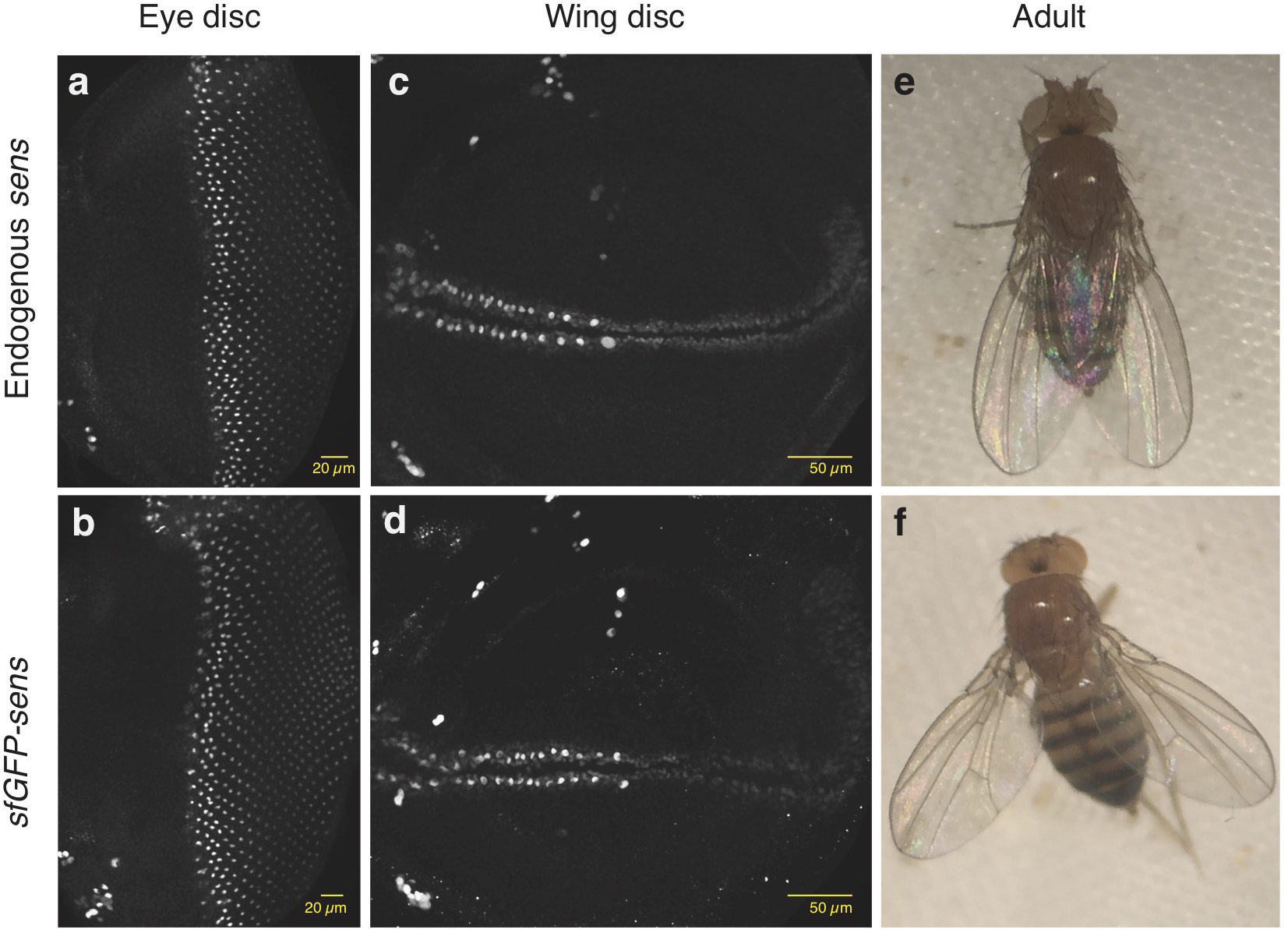
The *sens* transgene rescues endogenous Sens expression and null mutant phenotypes. **a-d,** Imaginal discs stained with anti-Sens. Endogenous *sens* protein expression in **a,** wildtype eye and **c,** wing discs. *sfGFP-sens* protein expression in **b,** eye and **d,** wing discs. No endogenous protein is present in these discs, **e-f,** Adults with two copies of the sens transgene **f,** show no phenotypic differences compared to wild type adults **e.**

**Extended Data Figure 2.**
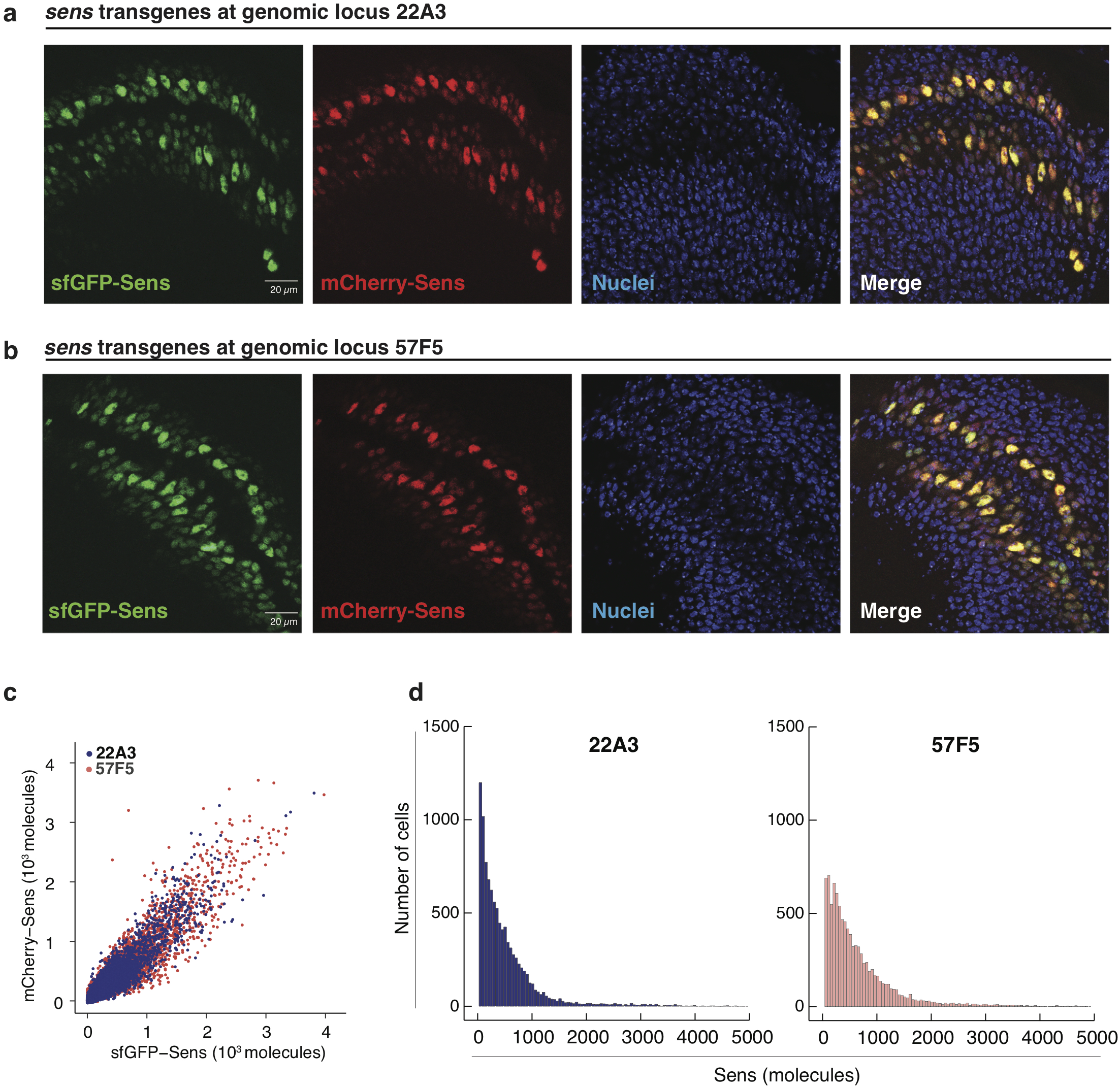
*sens* transgenes inserted at 22A3 and 57F5 landing sites show similar Sens expression patterns and protein abundance. **a,** sfGFP- and mCherry-tagged Sens in white prepupal wing discs expressing sens inserted at landing site 22A3 or **b,** 57F5. Nuclei are stained with DAPI (blue), **c,** Scatter plot of single-cell Sens protein levels from sens alleles inserted at 22A3 or 57F5. **d,** Flistograms depicting the distribution of Sens-positive cells sorted by Sens abundance. 10,000 cells were randomly sampled from 22A3 and 57F5 datasets each for comparison.

**Extended Data Figure 3.**
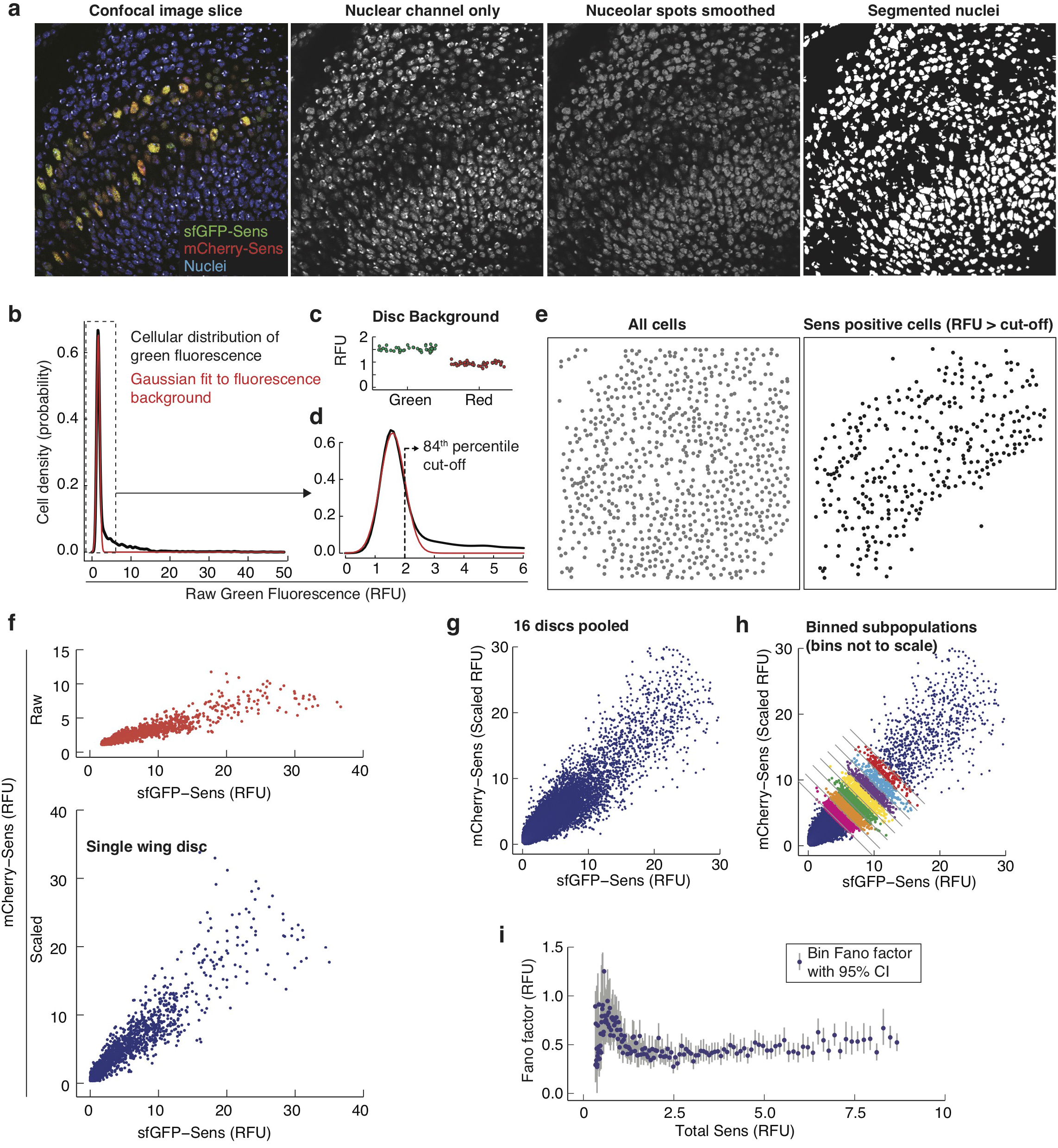
Image analysis and Fano factor calculation from nuclear fluorescence signals of Sens positive cells. **a,** The nuclear (DAPI) channel from individual confocal image slices was used to computationally identify and segment single nuclei. Average single cell sfGFP and mCherry fluorescence signals were then estimated from segmented nuclei, **b,** Raw sfGFP signal (green fluorescence channel) histogram from the wing disc in **a,** is shown as an example. The Gaussian-fltted fluorescence background is shown in red. **c,** Mean fluorescence background was calculated for green or red channels in individual images. Each datapoint is one disc. **d,** Magnified view of histogram in **b.** Dashed line marks the cut-off signal value above which cells were considered Sens-positive, e. Computationally identified Sens-positive cells displayed the expected expression pattern and were used for further analysis. **f,** The mCherry signal intensity is scaled relative to sfGFP signal in Sens-positive cells from individual discs. **g,** Data from discs of the same genotype are pooled together. **h,** Pooled data is sorted into separate bins according to total Sens signal. The variance and mean Sens signal is calculated for each binned sub-population. **i,** The Fano factor (variance/mean) of each bin is plotted as a function of mean Sens signal in a bin. 95% confidence intervals for the estimated Fano factor are calculated by boot-strapping with resampling within each binned subpopulations.

**Extended Data Figure 4.**
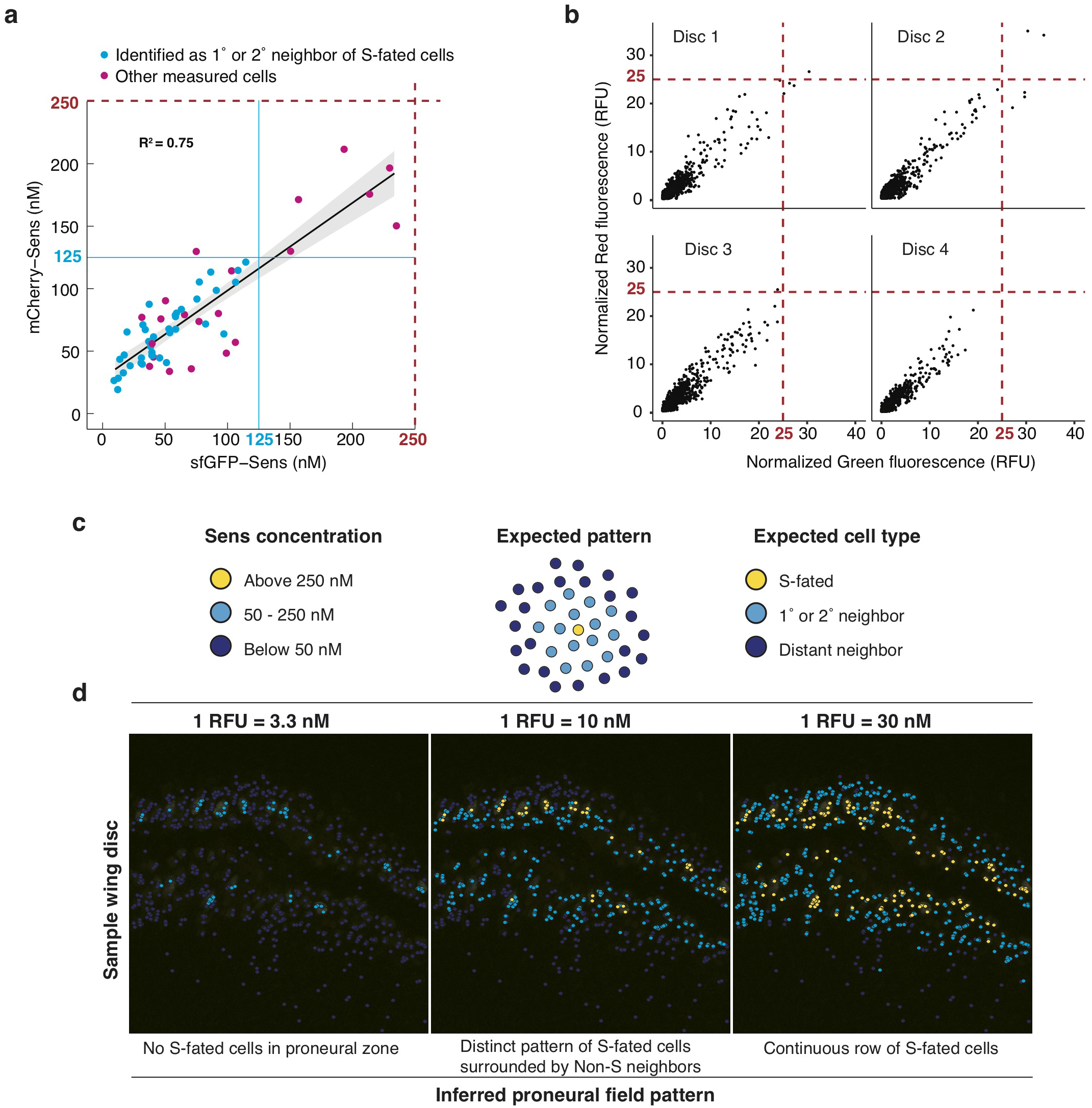
Comparison of Fluorescence Correlation Spectroscopy (FCS) and confocal microscopy fluorescence measurements to convert Relative Fluorescence Units (RFU) into absolute concentration of Sens (nM). **a,** FCS measurements of mCherry-Sens and sfGFP-Sens concentrations in live wing disc nuclei. Cyan cells were directly identified as first degree (1°) or second degree (2°) neighbors of S-fated cells. Protein output from both alleles is positively correlated (linear regression fit in black with 95% confidence of fit in grey) with almost no significant difference in output between the alleles. Neighbors of S-fated cells express Sens in the range 25 - 125 nM protein per allele (cyan lines). **b,** Relative fluorescence intensity (RFU) for cells in four fixed wing discs is shown. Red dashed lines mark the estimated maxima for Sens expression at 25 RFU, corresponding to 250 nM maxima measured by FCS. **c,** Expected arrangement of S-fated cells (yellow), first or second degree neighbors (cyan), and distant neighbors (dark blue) within Sens-positive stripes adjacent to the wing margin. Cells were categorized into distinct classes based on total Sens concentration measured by FCS (left). **d,** Fluorescence signal intensity (RFU) from imaged cells was converted into concentration (nM) by multiplying with a particular conversion factor. Cell coordinates were then mapped in space, and color-coded according to their expected cell category (defined in **c)**. Three conversion factors were tested, as shown at top. The factors differed from one another across a nine-fold range. Only a conversion factor of 10 nM/RFU (center column) was able to recapitulate the expected pattern of S-fated cells, as shown in the sample wing disc. Decreasing the factor to 3.3 (left) eliminated S-fated cells. Increasing the factor to 30 (right) produced a continuous row of S-fated cells. Thus, we estimate the conversion factor to be within an order of magnitude of the estimated conversion of 1 RFU equivalent to 10 nM.

**Extended Data Figure 5.**
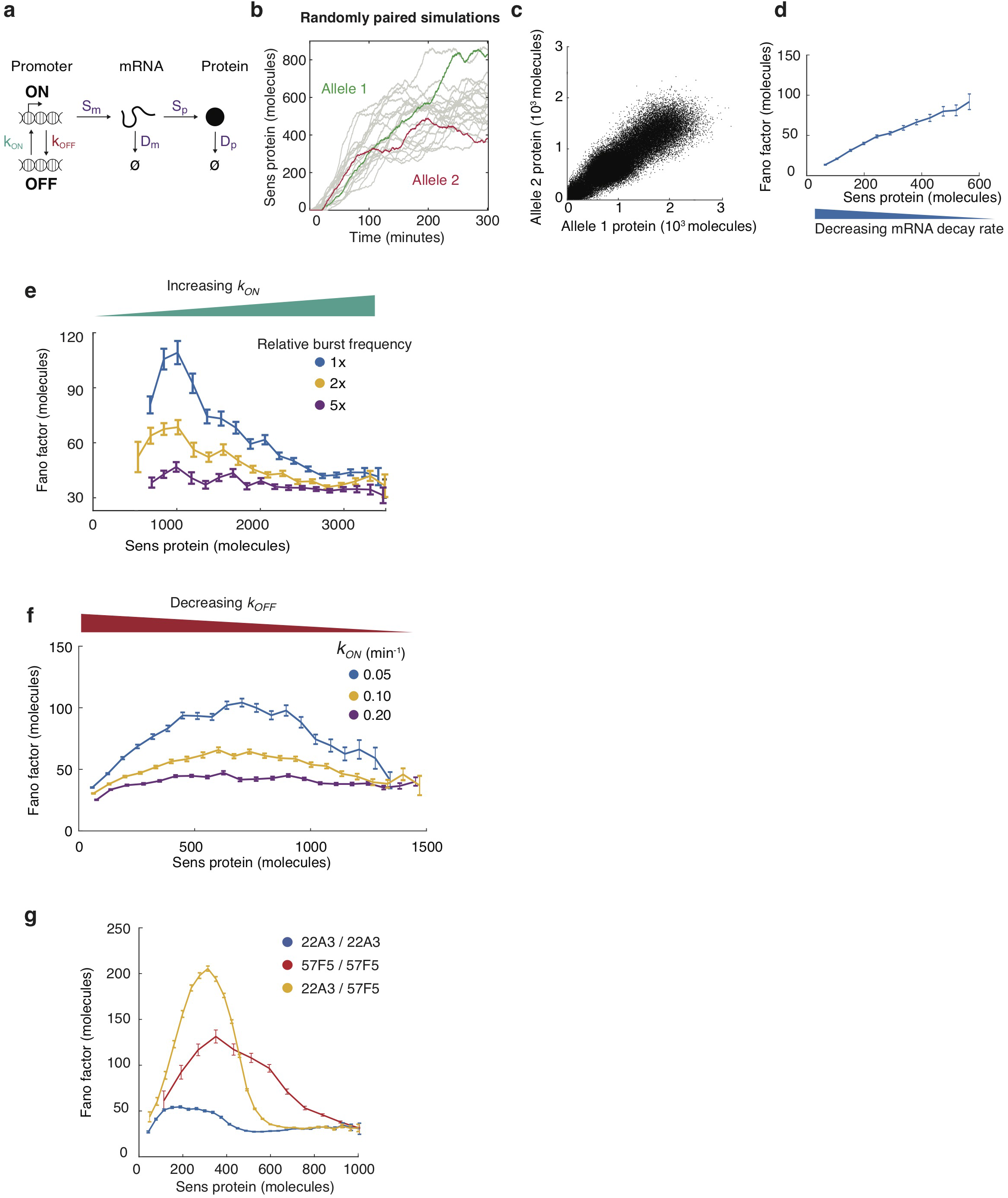
Modeling of Sens expression and noise. **a,** Simple model of gene expression with six rate parameters as shown. **b,** Simulated trajectories of Sens protein molecule numbers over time starting from zero molecules. Two randomly paired trajectories are shown in green and red, which mimic two independently acting alleles within the same cell, **c,** Sens protein output from randomly paired virtual alleles. Sampling was taken when all simulations had reached steady state. The *k_on_* rate parameter was varied to generate a range of steady-state protein output, and pairing was only done between simulations with identical rate parameters. The full range of Sens expression can be recapitulated by varying any of the six gene expression rate parameters in the model, **d-f.** Simulations were performed by systematically varying a particular rate parameter, and the Fano factor was calculated for bins of randomly paired simulations according to their protein output. Error bars are 95% confidence intervals as calculated by bootstrapping, **d,** The mRNA decay rate constant *D_m_* is systematically varied to generate a range of Sens expression. The resulting Fano factor slowly rises as protein levels increase. Similar profiles were obtained for gradients created by rate parameters *S_p_* and *D_p_*. **e,** Promoter switch ON rate *k_on_* is systematically varied to generate a range of Sens expression. Within this regime, burst frequency (*k_on_ · k_off_/k_on_ + k_off_*) is determined by both rates *k_on_* and *k_off_*. Shown are Fano profiles when promoter switching rate per unit time is increased by proportionately increasing both *k_on_* and *k_off_* 2 or 5-fold, such that the ratio *k_on_/k_off_* remains constant. As shown, when promoter switching is faster, the Fano peak diminishes because subsequent transcription bursts become more frequent, **f,** *k_off_* is systematically varied to generate a range of Sens expression. *k_off_* impacts both burst size (*S_m_/k_off_*) and burst frequency (*k_on_ · k_off_/k_on_ + k_off_*). Greatest noise is observed at approximately half maximal protein output. Shown are Fano profiles when *k_on_* is fixed to one of three different values. Again, the model suggests that when burst frequency alone is increased (by increasing *k_on_* in this case), the Fano factor is reduced. **g,** Fano profiles from simulations in which paired alleles are assumed to act independent of one another. One panel of simulations had the burst size (*S_m_/k_off_*) set to 5 mRNAs/burst mimicking 22A3, and another panel’s burst size was set to 10 mRNAs/burst mimicking 57F5. Random pairing of the 22A3 and 57F5 simulations generates a large Fano peak amplitude. Error bars are 95% confidence intervals.

**Extended Data Figure 6.**
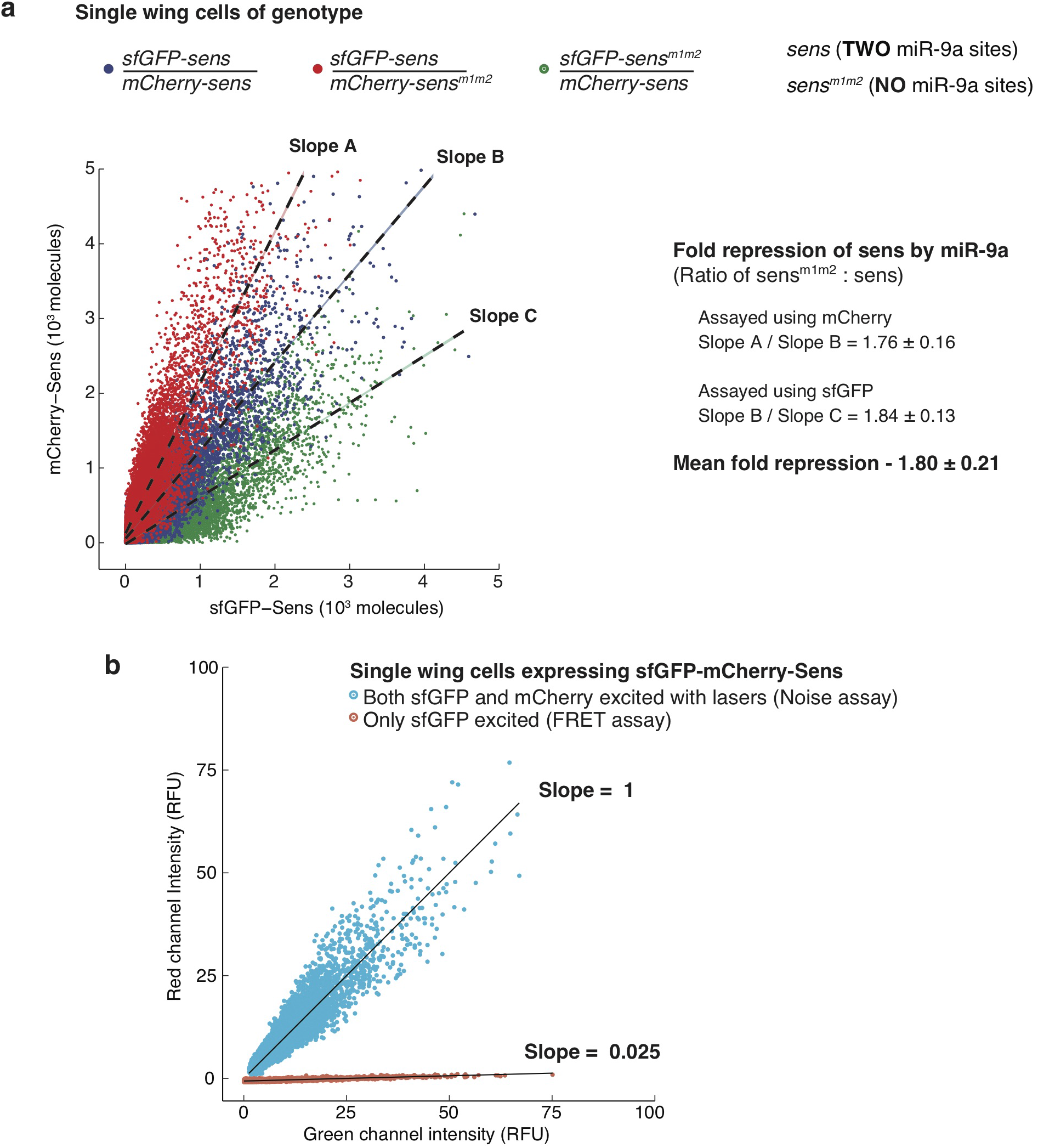
Fluorescent protein tags sfGFP and mCherry behave interchangeably *in vivo* and display negligible energy transfer under experimental imaging conditions. **a,** Single wing disc cells of the indicated genotypes were imaged and their nuclear sfGFP and mCherry fluorescence intensity was measured. Cells from individual discs were pooled separately to calculate the mCherry:sfGFP linear slope. Slopes were compared across genotypes (with relative errors propagated) to sensitively assay if the nature of the fluorescent protein tag interferes with quantitative protein measurements. Red dots represent single cells containing a *mCherry-sens* allele free of miR-9a control and an *sfGFP-sens* allele under miR-9a regulation. Blue dots represent cells where both alleles are under miR-9a repression. Green dots represent single cells containing an *sfGFP-sens* allele free of miR-9a control and a *mCherry-sens* allele under miR-9a regulation. Sens protein output was measured in cells with or without miR-9a regulation. As seen, mCherry-Sens protein is consistently lower in blue cells relative to their red counterparts, across the entire range of Sens expression. The ratio of slopes for the two groups provides the fold reduction in Sens protein due to miR-9a activity. Similarly, sfGFP-Sens protein is consistently lower in blue cells relative to their green counterparts. Assaying miR-9a activity using either fluorescent tag shows no significant difference such that on average miR-9a decreases protein output 1.8-fold across the entire range of Sens expression. Error bars represent standard error of the mean. **b,** Fluorescence Resonance Energy Transfer (FRET) from sfGFP to mCherry molecules is negligible under experimental imaging conditions. Wing disc cells with tandem tagged *sfGFP-mCherry-sens* alleles were imaged under identical conditions with either both sfGFP and mCherry molecules excited by lasers to assay stochastic noise, or with only sfGFP excited to assay FRET from sfGFP to mCherry molecules. Raw single cell green channel fluorescence intensity is plotted on the x-axis. All cells in the imaging field were included for analysis. Red channel fluorescence values, on y-axis, were linearly transformed to make the slope equal to 1 and y-intercept equal to 0 for noise assay data. Single cell red fluorescence values for FRET assay data were then transformed with identical parameters for comparision.

**Extended Data Figure 7.**
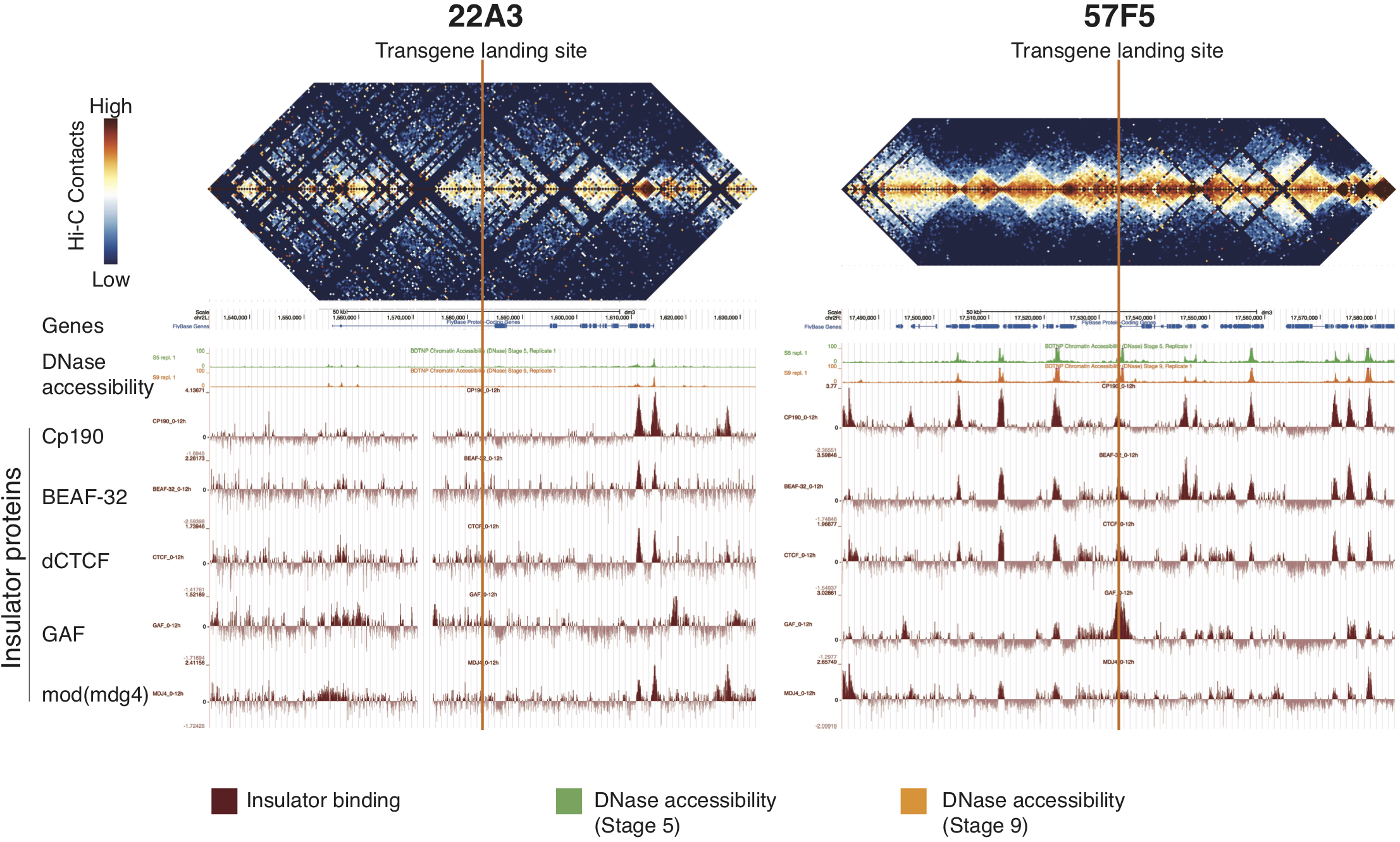
*sens* transgene landing sites 22A3 and 57F5 differ in chromatin compaction and distance from insulator protein enriched boundaries of Topologically Associated Domains (TADs). Top: Heat maps of aggregate Hi-C data used to calculate chromosomal contact frequency for embryonic ncl4 datasets (from Stadler et al., 2017)^27^ is shown for 100 kb windows centered on the landing sites at 22A3 (left) and 57F5 (right). Vertical orange lines denote the precise locations of the two transgenic landing sites. Bottom: A UCSC browser window for the corresponding coordinates is shown with tracks for annotated genes, DNase accessibility (from Li et al., 2011)^39^, and ChlP-seq of the insulator proteins CP190, BEAF-32, dCTCF, GAF and mod(mdg4) (from Nègre et al., 2010)^40^. Although these data were derived from embryonic genomes, Stadler et al., (2017)^27^ showed that the embryonic TAD boundary elements correspond to the locations of mapped interband regions of third-instar larval polytene chromosomes. This strongly suggests that TAD organization is largely maintained during the development of embryos into third-instar larvae. Thus, it is highly likely that the genome organization at 22A3 and 57F5 in larval wing disc cells is similar to this mapped data.

**Extended Data Figure 8.**
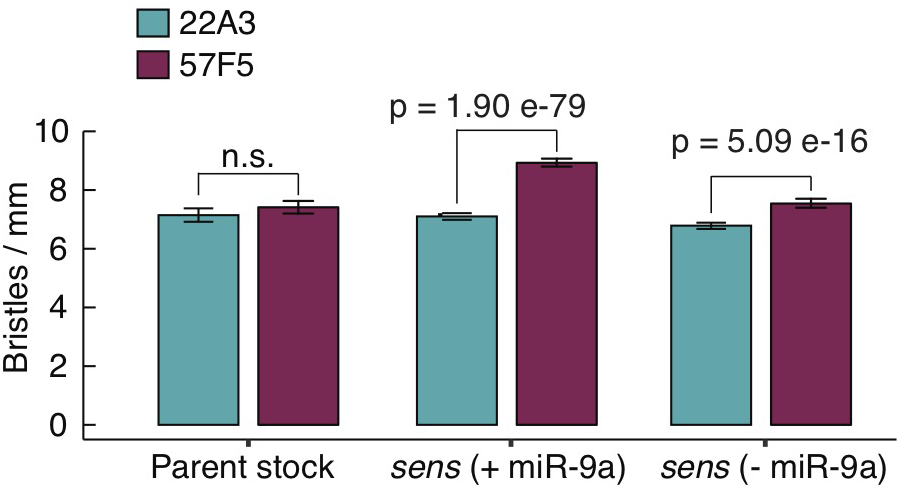
Genomic location of *sens* increases the density of chemosensory bristles specified in adults. If ectopic chemosensory bristles arise due to Sens protein fluctuations, they would decrease the average distance between adjacent chemosensory bristles. Therefore, we measured chemosensory bristle density on the anterior wing margin of adult females. When *sens* is expressed from 57F5, chemosensory bristle density is greater relative to 22A3. This effect is observed irrespective of the nature of *sens* allele compared i.e. with or without miR-9a regulation of *sens*. In contrast, bristle density is not significantly different between the parental 22A3 and 57F5 stocks which express *sens* from the endogenous locus on chromosome III. Error bars are 95% confidence intervals and p values are from a student’s t-test.

**Extended Data Figure 9.**
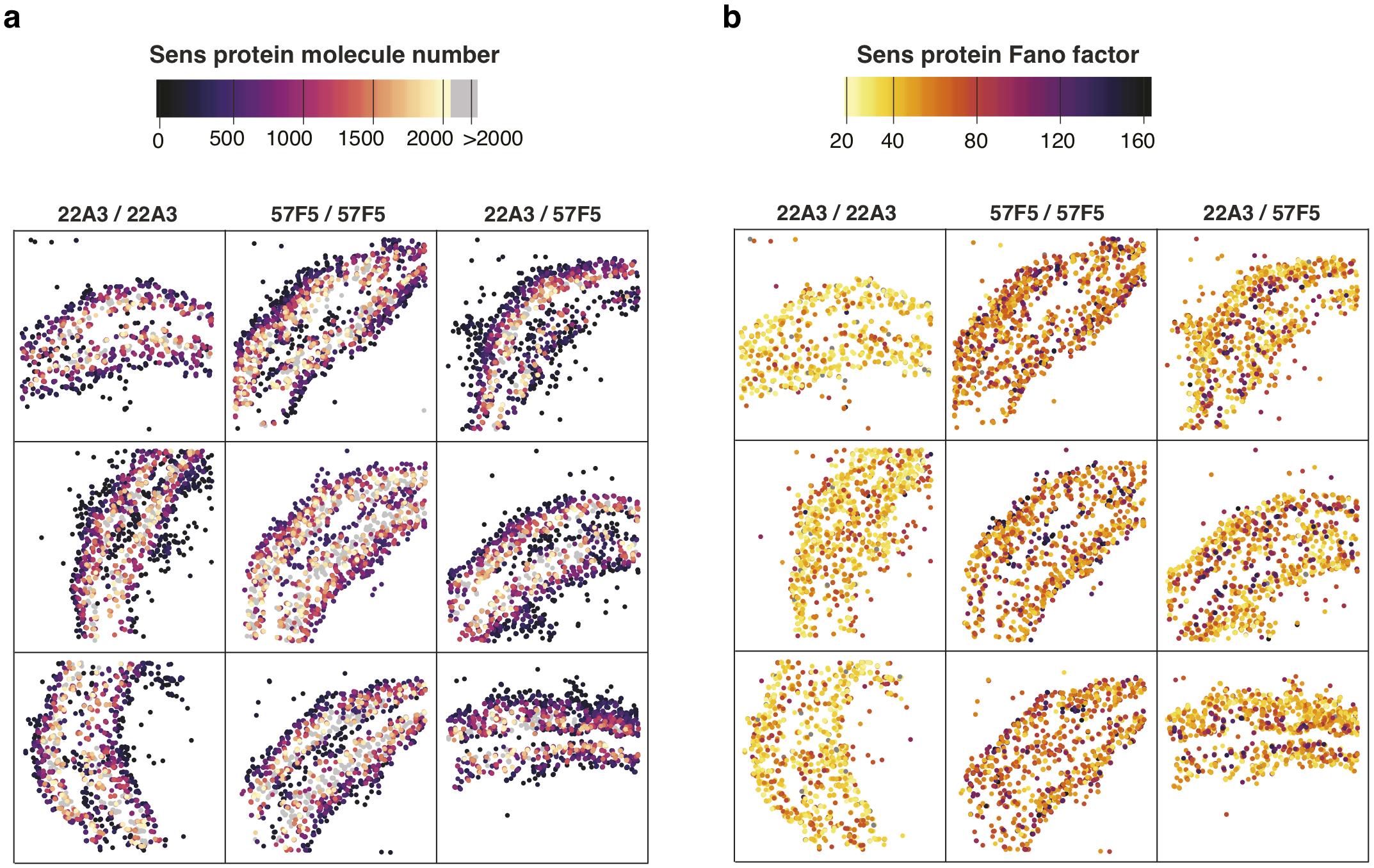
Fano factor, but not abundance, of Sens protein is dependent on genomic location of the *sens* gene. Spatial map of cell coordinates from individual wing discs are shown. **a-b,** Representative wing discs with homologous allele pairs for sens at 22A3 (left), 57F5 (center) and the non-homologous pair 22A3/57F5 (right) are shown. Cell centroids are color coded according to their corresponding **a,** Sens protein abundance or **b,** Fano factor. Color scale on top. Only cells with fewer than 2000 molecules of Sens are plotted for ease of visualization.

## Methods

### Generation of sens transgenic stocks

The sens alleles *sens^E1^* and *sens^E2^* are protein null mutants^9,35^. N-terminal 3xFlag-TEV-StrepII-sfGFP-FlAsH tagged sens originally generated from the CH322-01N16 BAC was a kind gift from K. Venken and H. Bellen^14^ and has been shown to rescue *sens^E1^* and *sens^E2^* mutations^14,15^. To generate mCherry tagged *sens*, the sfGFP coding sequence in 3xFlag-TEV-StrepII-sfGFP-FlAsh was swapped out for mCherry by RpsL-Neo counter-selection (GeneBridges). The sfGFP-mCherry tandem tagged *sens* transgene was generated similarly by overlap PCR such that sfGFP and mCherry sequences were separated by a 12 amino acid (GGS)_4_ linker. The *miR-9a* binding site mutant alleles of the tagged sens transgenes were created by deletion of the two identified binding sites in the 607 nt sens 3’ UTR as had been described previously^15^ to generate *sens^m1m2^* mutant transgenes. Cloning details are available on request. All BACs were integrated at PBacy+-attP-3BVK00037 (22A3) and PBacy+-attP-9AVK00022 (57F5) landing sites by phiC31 recombination^14^. Transgenic lines were crossed with sens mutant lines to construct stocks in which sens transgenes were present in a *sens^E1^/sens^E2^* trans-heterozygous mutant background. Thus, the only Sens protein expressed from these animals came from the transgenes. All experiments were performed on these stocks.

### Adult wing imaging and mispatterning analysis

Adult females from uncrowded vials were collected on eclosion and aged for 1-2 days before being preserved in 70% Ethanol. Wings from preserved animals were plucked out with forceps and kept ventral side up on a glass slide. Approximately 10 pairs of wings were arranged per slide using a thin film of Ethanol to lay them flat. Left and right wings from the same animal were positioned next to each other. Once specimens were arranged as desired, excess ethanol was wiped away. A second glass slide was coated with heptane glue (10 cm^2^ double sided embryo tape dissolved overnight in 4 ml heptane) and pressed down onto the specimen slide to affix them dorsal side up. Then wings were mounted in 70% glycerol in PBS and sealed with nail polish for imaging. Wings were imaged using an Olympus BX53 upright microscope with a 10x UPlanFL N objective in brightfield. To achieve optimal resolution, 8-10 overlapping images were taken for each wing and stitched together in Adobe Photoshop^®^.

Wings with at least one mechanosensory bristle placed ectopically in or adjacent to the chemosensory bristle row were counted as mispatterned. The proportion of mispatterned wings was calculated for each genotype (n ≥ 60). Genotypes were compared by calculating the odds ratio of mispatterning and determined to be significantly different from 1 if p < 0.05 using Fischer’s exact test. For chemosensory bristle density, wing images were used to identify and mark chemosensory bristles along the margin in Fiji. The euclidean distance between successive bristles was measured and bristle density was calculated as the inverse of mean spacing. 95% confidence intervals were calculated by bootstrapping and bristle distributions across genotypes were compared statistically using student’s t-test.

### Fluorescence microscopy

All fluorescence microscopy experiments used female white pre-pupal animals. The white pre-pupal stage was chosen because it is easily identified and lasts for only 45-60 minutes. Further, wing margin chemosensory precursor selection was observed to be tightly linked to the transition from late third larval instar to pre-pupal stage. Therefore,choosing white pre-pupal animals allowed us to strictly control for developmental stage. Wing discs from staged animals were dissected out in ice-cold Phosphate Buffered Saline (PBS). Discs were fixed in 4% paraformaldehyde in PBS for 20 minutes at 25°C and washed with PBS containing 0.3% Tween-20. Then they were stained with 0.5 *μ*g/ml DAPI and mounted in Vectasheild™. Discs were mounted apical side up and imaged with identical settings using a Leica TCS SP5 confocal microscope. All images were acquired at 100x magnification at 2048 x 2048 resolution with a 75 nm x-y pixel size and 0.42 *μ*m z separation. Scans were collected bidirectionally at 400 MHz and 6x line averaged. Wing discs of different genotypes were mounted on the same microscope slide and imaged in the same session for consistency in data quality.

### Immunohistochemistry

Discs were dissected and fixed as above before incubating with the primary antibodies of interest. Tissues were washed thrice for 5-10 minutes each and incubated with the appropriate fluorescent secondary antibodies (diluted 1:250) for 1 hour. They were then stained with DAPI, washed in PBS-Tween and mounted for imaging. Guinea pig anti-Sens antibody (gift from H. Bellen) was used at 1:1000 dilution.

### Image analysis

#### Cell Segmentation

For each wing disc, five optical slices containing proneural cells were chosen for imaging and analysis. A previously documented custom MATLAB script was used to segment nuclei in each slice of the DAPI channel^36,37^. Briefly, high intensity nucleolar spots were smoothed out to merge with the nuclear area to prevent spurious segmentation. Next, cell nuclei were identified by thresholding based on DAPI channel intensity. Segmentation parameters were optimized to obtain nuclei with at least 100 pixels and no more than 4000 pixels. For each nuclear area so identified, the average signal intensity for the sfGFP and mCherry channels was recorded along with the relative position of its centroid in x and y. Since segmentation was based exclusively on the nuclear signal, it identified all cells present in the imaged area (Extended Data Fig. 3a).

#### Background fluorescence normalization

The majority of cells imaged did not fall within the proneural region and therefore displayed background levels of fluorescence scattered around some mean level (Extended Data Fig. 3b). Sens expressing cells were present in the right hand tail of the distribution. The background was channel specific and varied slightly from disc to disc (Extended Data Fig. 3c). Therefore, we calculated the mean channel background for each channel in each disc individually. We did this by fitting a Gaussian distribution to the population and finding the mean of that fit. In order to separate Sens positive cells, we chose a cut-off percentile based on the normal distribution, below which cells were deemed Sens negative. We set this cut-off at the 84^th^ percentile for all analysis (Extended Data Fig. 3d).

This was determined empirically by mapping cell positions relative to the pronueral region. At and above the 84^th^ percentile, mapped cells followed the proneural striped pattern. Lowering the cut-off led to addition of cells randomly scattered across the imaging field. Increasing the cut-off led to progressive narrowing of the proneural stripes. From this we inferred the fluorescence level at 84^th^ percentile as a tolerant but specific threshold to identify Sens positive cells. Thus, to normalize measurements across tissues and experiments, this value was subtracted from the total measured fluorescence for all cells in that disc and channel. Only cells with values above the threshold for both mCherry and sfGFP fluorescence were assumed Sens positive (usually 30% of total cells) and carried forward for further analysis (Extended Data Fig. 3e).

#### mCherry and sfGFP fluorescence units scaling

We required the relative fluorescence of the mCherry and sfGFP channels to be scaled in equivalent units. To do this, we fit a linear equation as shown, and derived best-fit values for slope and constant intercept.

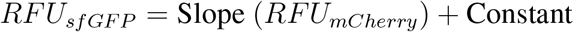

To preserve data integrity, the slope and constant was calculated for each wing disc separately. Linear correlation coefficients were consistenly high between mCherry and sfGFP fluorescence, ranging from 0.85 to 0.95. Finally, to rescale single cell mCherry fluorescence in units of sfGFP-Sens fluorescence, we applied the following transformation to each cell’s mCherry reading (Extended Data Fig. 3f).

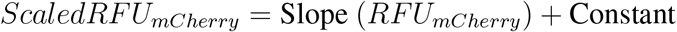

Once the two-channel RFUs were made equivalent, they were summed to obtain total Sens RFU for each cell as shown.

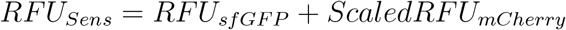

#### Stochastic noise and Fano factor calculation according to gene expression level

We used the following formula to calculate intrinsic noise^1^. Mathematically, it is the variance remaining after the co-variance term of two variables is subtracted from their total variance. This value is then normalized to the squared mean (*η*^2^ = *σ*^2^/*μ*^2^) to obtain the following dimensionless quantity:

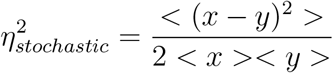

Here *x* and *y* represent *RFU_sfGFP_* and *ScaledRFU_mcherry_*. Angled brackets denote averages over the cell population. This term provides a single value of intrinsic noise for the entire cell population. Since Sens expression varies over three orders of magnitude, we partitioned cells into smaller bins according to their total Sens expression. We then calculated intrinsic noise for each binned sub-population. Sens expression RFU was log-transformed and we used a bin width of 0.02 log(RFU) to partition cells (Extended Data Fig. 3g-h).

For each bin, we calculated the intrinsic noise *η*^2^ as well as mean Sens expression *μ*. These were multiplied together to calculate the Fano factor for each bin.

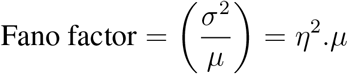

Given that the number of cells in each bin was not constant, and that variance estimates are affected by sample size, we calculated confidence intervals around the calculated Fano factor for each bin by bootstrapping. We resampled bin populations 50,000 times with replacement. The 2.5^th^ and 97.5^th^ percentile estimates were used to construct a 95% confidence interval for that bin’s Fano factor (Extended Data Fig. 3i).

#### Technical noise subtraction

Intrinsic noise and the Fano factor were calculated as described above for tandem-tagged *sfGFP-mCherry-sens* wing discs. Intrinsic noise was identical for tandem-tagged *sens* genes inserted at either 22A3 or 57F5. This would be expected if the intrinsic noise from this transgene was caused by stochastic processes related to photon detection and counting. Therefore, we pooled data generated from both locations before binning into sub-populations. In order to construct a statistical model for technical noise at each level of Sens fluorescence, we used a Lowess regression to fit a continuous line through the data (as seen in Fig. 2a). The Lowess algorithm fits a locally weighted polynomial onto x-y scatter data and therefore does not rely upon specific assumptions about the data itself. The local window used to calculate a fit was kept constant for all Lowess fits. Using our statistical model, we generated a predicted Fano factor that was due to technical noise for each bin. This predicted value was subtracted from the Fano factor that was due to both technical and gene expression noise for each bin. The difference obtained is an estimate of the Fano factor due to noise in *sens* gene expression.

#### Fluorescent tag similarity

As an additional control, we checked by various means if indeed sfGFP-Sens and mCherry-Sens proteins behaved similarly *in vivo* such that the nature of the protein tag did not affect quantitative assays.

First, we measured the molecule counts of sfGFP-Sens and mCherry-Sens in the same cells using Fluorescence Correlation Spectroscopy (FCS). As can be seen in Extended Data Fig. 4a, we obtained similar numbers of Sens molecules irrespective of which fluorescent tag was attached. This indicated that both alleles express equal numbers of protein *in vivo*.

Second, using the microRNA repression assay detailed in Extended Data Fig. 6a, we sensitively assayed whether the nature of the tag affects protein output quantitatively. If one tag were differentially expressed relative to the other, we would expect the fold-repression values calculated using mCherry tagged *sens* alleles to be different from sfGFP tagged *sens* alleles. This was not observed.

Third, to ensure that we did not under-sample stochastic noise due to Fluorescence Resonance Energy Transfer (FRET), we imaged tandem tagged sfGFP-mCherry-Sens samples in both channels after exciting only the donor (sfGFP) molecules. As shown in Extended Data Fig. 6b, there is negligible FRET from sfGFP to mCherry when using imaging parameters identical to experimental runs.

### Fluorescence Correlation Spectroscopy (FCS)

#### FCS sample calibration and measurement

White pre-pupal wing discs were dissected in PBS and sunken into LabTek 8-well chambered slides containing 400 *μ*l PBS per well^38^. Discs were positioned such that the pouch region was facing the bottom of the well to be imaged. FCS measurements were made using an inverted Zeiss LSM780, Confocor 3 instrument with APD detectors. A water immersion 40x objective with numerical aperture of 1.2 (which is optimal for FCS measurements) was used throughout. Fast image scanning was utilized for identification of cell nuclei to be measured by FCS. Prior to each session, we used 10 nM dilute solutions of Alexa488 and CF586 dyes to calculate the average number of particles, the diffusion time and define the structural parameters 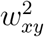 and *z*_0_. Using these we calibrated the Observation Volume Element (OVE) whose volume can approximated by a prolate ellipsoid 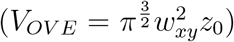. Measurements were performed in Sensory Organ Precursor cells (SOPs or S-fated), as well as first and second order neighbors, residing dorsally or ventrally of the S-fated cell (Extended Data Fig. 4a). Measurements were subjected to analysis and fitting, using a two components model for three dimensional diffusion and triplet correction as follows:

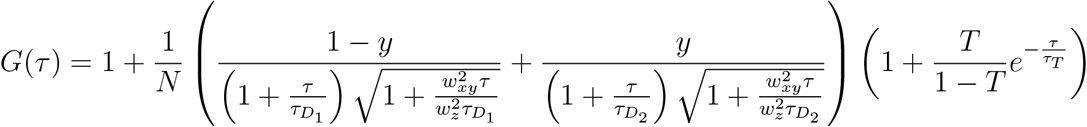

FCS measurements were excluded from analysis if they exhibited marked photobleaching or low CPM i.e. counts per molecule (CPM < 0.5 kHz per molecule per second). Due to the higher CPM of sfGFP, it was expected that Sens-sfGFP measurements are more accurate. We, nevertheless, observed fairly similar molecular numbers for both sfGFP-Sens and mCherry-Sens. Normalized auto correlation curves allowed us to compare the differential mobilities of the tagged Sens protein molecules in the nucleus and their degree of interaction with chromatin. Consistently, for both sfGFP and mCherry tagged transcription factors, we observed similar amplitudes and decay times of the slow FCS component, suggesting that the interaction with chromatin is not substantially different for differently tagged Sens molecules or even at different Sens concentrations.

#### Conversion from relative fluorescence to molecule counts

We compared Sens protein concentrations as measured by FCS to single cell fluorescence data from confocal imaging of the fixed tissue. All comparisons were done for the genotype shown below since all FCS measurements were made in these animals.

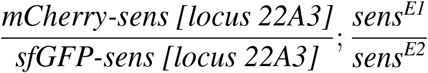

First, we looked at the extremes of Sens expression. Since FCS was only performed on Sens positive nuclei as visible by eye, we did not consider the lowest Sens expressing cells comparable to the confocal measured minimum Sens. However, we expected cells with highest Sens to be of similar magnitude between the two methods. To mitigate the effect of extreme outliers on the maxima, we looked at Sens expression profiles of individual discs (Extended Data Fig. 4b). As can be seen from FCS data, for both sfGFP and mCherry channel measurements, the highest Sens levels are no greater than 250 nM (per channel). The highest Sens positive cells, as measured by fluorescence microscopy, display approximately 25 RFU Sens (per channel). This gave us a rough conversion factor of 1 RFU equivalent to 10 nM.

Next we looked at the first and second order neighbors of S-fated cells. While S-fated cells show a large range of Sens expression, their E-fated neighbors display relatively less dispersion. FCS analysis showed that most of these cells expressed Sens in the range of 25nM - 125 nM per channel. Summing up the signal from sfGFP-Sens and mCherry-Sens, this corresponds to a total nuclear concentration of 50-250 nM. Based on this we divided Sens positive nuclei into three categories as shown in Supplementary Table 5.

We then mapped the labelled cell types to the original images. We expected that categorizing cells and mapping their positions should recreate the wing margin pattern i.e. S-fated cells (yellow) dispersed periodically along both sides of the wing margin surrounded by 1° and 2° neighbors (cyan) (Extended Data Fig. 4c). Indeed, we observe this pattern reproducibly if 1 RFU is assumed equivalent to 10 nM Sens (center column Extended Data Fig. 4d).

To further test this conversion factor, we made an order of magnitude estimation. Assuming 1 RFU = 3.3 nM (left column) or 1 RFU = 30 nM (right column), we again labelled cells according to the FCS observed cell types for different concentrations of Sens. As shown, increasing or decreasing the conversion factor three-fold does not reproduce the expected spatial pattern. This is most notable in the S-fated category where we see either none (3.3 nM) or a near-continuous stripe (30 nM) (Extended Data Fig. 4d).

Thus we chose 10 nM as a reasonable conversion factor from fluorescence to nanomolars for our data. Next, we converted from nanomolars to molecule numbers. Assuming a measured average wing disc cell nuclear volume of 22.99 x 10^−15^ L, each nanomolar of Sens corresponds to 13.8 molecules^38^. Therefore, we converted relative fluorescence units to molecules per nucleus as follows:

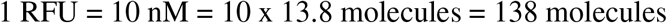

### miR-9a repression measurements

In order to measure the fold-decrease in Sens protein output due to miR-9a repression of *sens* mRNA, we compared the ratio of mCherry-Sens to sfGFP-Sens in the following genotypes:

A. Only *mCherry-sens* resistant to miR-9a repression

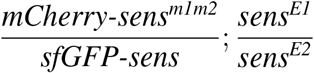
B. Neither *mCherry-sens* or *sfGFP-sens* resistant to miR-9a repression

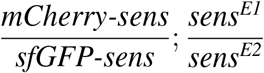
C. Only *sfGFP-sens* resistant to miR-9a repression

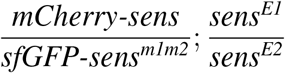

Single cell fluorescence values were obtained after cell segmentation and background subtraction as described earlier. Cells from individual discs were pooled together and red-green fluorescence was linearly correlated using least squares fit (QR factorization) to determine a slope and intercept for each disc. Next the average slope was calculated for each genotype (shown above).

Fold reduction in mCherry-Sens protein output due to miR-9a was calculated as the ratio of slope-(A) to slope-(B) with relative errors propagated. Similarly, fold reduction in sfGFP-Sens protein output due to miR-9a was calculated as the ratio of slope-(B) to slope-(C)(Extended Data Fig. 6a).

### Topological Domain Structure

Heat maps of aggregate Hi-C data were used to calculate chromosomal contact frequency for embryonic nc14 datasets^27^ (Extended Data Fig. 7, data from Stadler et al., 2017) for landing sites at 22A3 and 57F5. DNase accessibility data^39^ (from Li et al., 2011), and ChIP-seq of the insulator proteins CP190, BEAF-32, dCTCF, GAF and mod(mdg4) (from Négre et al., 2010)^40^ for the corresponding coordinates is shown as well.

### Experimental estimation of rate constants

#### mRNA decay rate *D_m_*

Female pre-pupal wing discs were dissected in WM1 medium^41^ at room temperature. To inhibit RNA synthesis, discs were incubated in WM1 plus 5 *μ*g/ml Actinomycin D in light protected 24-well dishes at room temperature. Approximately 20 discs were collected at 0, 10, 20 and 30 minutes post-treatment and were homogenized with 300 *μ*l Trizol for RNA extraction and qPCR analysis. Long-lived Rpl21 mRNA was used to normalize mRNA levels across time points. Similar results were obtained when 18S rRNA was used for normalization. mRNA decay was assumed exponential and a curve fit across all time-points was used to calculate the decay constant *D_m_* to be 0.0462 mRNA/min corresponding to a half-life *t*_1/2_ = 15.75 minutes (R^2^ = 0.91). Hsp70 mRNA decay was also measured as an additional short-lived control with known half-life (observed *t*_1/2_ = 35 mins). All qPCR primers used are listed in Supplementary Table 6

#### Protein decay rate *D_p_*

Homozygous *3xFlag-TEV-StrepII-sfGFP-FlAsH-sens* (in a sens null background) female pre-pupal wing discs were dissected in WM1 medium at room temperature. Discs were incubated in WM1 plus 100 μg/ml cycloheximide for 0, 1, 2 and 3 hours at room temperature. Ten discs were harvested at each time-point and snap frozen in liquid nitrogen. To assay Sens protein abundance, we used an indirect sandwich ELISA (enzyme-linked immunosorbent assay) protocol as follows. Frozen discs were homogenized in 150 *μ*l PBS containing 1% Triton-X, centrifuged to remove crude particulate matter and then incubated with rabbit anti-GFP (1:5000) overnight at 4°C in anti-Flag antibody coated wells. Wells were washed with PBS with 0.2% Tween 20 and incubated with HRP linked goat anti-rabbit (1:5000) antibody for 2 hours at 37°C. Wells were subsequently washed and incubated with 100*μ*l 1-Step Ultra TMB-ELISA substrate. HRP activity was terminated after 30 minutes with 100*μ*l 2M H_2_SO_4_ and absorbance measured at 450 nm. Protein decay was assumed exponential and a curve fit across all time-points was used to estimate the decay constant *D_p_* to be 0.12 proteins/hr, corresponding to *t*_1/2_ = 5.09 hours (R^2^ = 0.84).

#### Protein synthesis rate *S_p_*

As has been theorized previously^17,18,23^ and also suggested by our experimental data (Fig.3), a constant Fano factor is related to the translation burst size *b* as follows

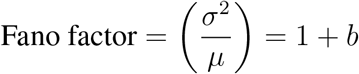

Here *b* is defined by the post-transcriptional rate constants as:

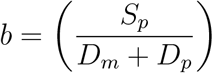

The Fano factor in the constant regime for Sens is ~20 molecules (Fig. 3c). Assuming *b* =19 and substituting the measured values for *D_m_* and *D_p_*, we estimate that *S_p_* ~ 0.9 proteins/mRNA/min. When miR-9a binding sites are deleted from the gene, Sens protein output is 1.80 ± 0.21 fold higher and the Fano factor is ~ 35 molecules (Fig. 3b-c). This makes the miR-9a resistant protein synthesis rate *S_p_* ~ 1.7 proteins/ mRNA/min. Thus, we fixed *S_p_* at 0.9 proteins/mRNA/min or at 1.7 proteins/mRNA/min to simulate *sens* alleles with and without miR-9a binding sites respectively.

### Stochastic Simulation Model

We modeled the various steps of gene expression, based on central dogma, as linear first order reactions (Extended Data Fig. 5a). To simulate the stochastic nature of reactions, we implemented the model as a Markov process using Gillespie’s stochastic simulation algorithm (SSA)^42^. A Markov process is a memoryless random process such that the next state is only dependent on the current state and not on past states. Simple Markov processes can be analyzed using a chemical master equation to provide a full probability distribution of states as they evolve through time. The master equation defining our three-variable gene expression Markov process is as follows:

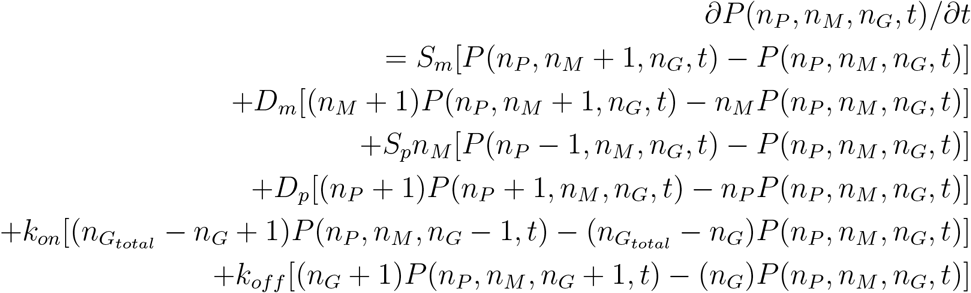

Here *n_P_* and *n_M_* denote the number of protein and mRNA molecules respectively. *n_G_total__* is the total number of genes of which *n_G_* are genes in the ‘ON’ state capable of transcription. Therefore, *n_G_/n_G_total__* is the fraction of active genes. Time is denoted by t. The rate constants are defined in Supplementary Table 2.

As the Markov process gets more complex, the master equation can become too complicated to solve. Gillespie’s SSA is a statistically exact method which generates a probability distribution identical to the solution of the corresponding master equation given that a large number of simulations are realized.

#### Simulation set-up and algorithm

The gene expression model is comprised of six events (Extended Data Fig. 5a) and their associated reaction rates shown in Supplementary Table 2. Unless specified, the events and rate constants were kept identical between *sfGFP-sens* and *mCherry-sens* alleles simulated in the same cell. At any given instance, for a given allele, either of these six events could take place.

Gillepsie’s SSA is based on the fact that the time interval between successive events can be drawn from an exponential distribution with mean 1/*r_total_* where

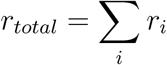

i.e the sum total of reaction rates for all *i* events. Further, the identity of the event that will occur is drawn from a point probability defined as

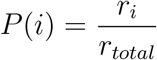

The algorithm proceeded as follows:

1. We initialized all simulations to start with state

Promoter state = off
mRNA molecules = 0
protein molecules = 0
simulation time = 0 minutes
2. *r_total_* was determined by calculating the individual rates *r_i_* at current time *t* which depend on the number of substrate molecules and the rate constants in Supplementary Table 2.
3. A random time interval *τ* was picked from the exponential distribution with mean 1/*r_total_*
4. A random event i was picked with probability *P*(*i*) as described above.
5. The cellular state was changed in accordance with the chosen event. The possible state changes were as follows

- Promoter state from off → on
- Promoter state from on → off
- mRNA molecule count increased by 1
- mRNA molecule count decreased by 1
- protein molecule count increased by 1
- protein molecule count decreased by 1
6. Simulation time was updated as *t* + *τ*
7. Steps 2 to 6 were iterated until total simulation time reached 5 hours.

#### Fano factor calculation

We ran simulations for 5 hours to approximate steady state expression, at the end of which protein and mRNA molecules produced from each allele in each cell were counted. A minimum of 5000 such ‘cells’ were simulated for each set of parameters. For simulations that tested the effect of parameter gradients on Sens noise, we divided the graded parameter into 20 discrete levels. Each level was simulated separately after which cells from all levels were pooled to generate a whole population. This population was binned into 25-30 bins based on total Sens level, and the Fano factor was calculated for each bin. Bootstrap with resampling was used to determine 95% confidence intervals for each bin’s Fano factor.

#### Parameter constraints

To keep simulations computationally feasible, we adjusted the slowest rate parameter, the protein decay rate *D_p_*, from 0.002 proteins/min to 0.01 proteins/min (half-life from 5 hours to 1 hour). This is because we conducted simulations until protein conditions reached steady state, which is approximately five-fold longer than the half-life for the slowest reaction. For 25-hour simulations, this was resource and time-intensive. We compared the noise trends in simulations with either *D_p_* of 0.002 proteins/min or to 0.01 proteins/min, and found both generated similar noise trends to one another. This indicates that protein decay is a not a major source of intrinsic noise in this model. Therefore, we kept *D_p_* at 0.01 proteins/min.

The transcriptional parameters *S_m_, k_on_* and *k_off_* were varied in accordance with the specific hypothesis being tested. We constrained them loosely to be within an order of magnitude of reported values for these rates from the literature^43^. We also constrained these rates so as to produce steady state protein numbers and Fano factors similar to experimental data. The minimum and maximum values used are listed in Supplementary Table 2.

#### Modeling *sens* regulation by a morphogen gradient

As seen in Fig. 1e, Sens-positive cells display a wide range of expression and they are patterned in space as stripes. This is due to signaling via Wg, which is secreted from the presumptive wing margin and diffuses to form a bidirectional morphogen gradient. Wg signaling directly activates transcription of the *sens* gene^8^. We assumed that at least one of the three transcriptional rate parameters (*S_m_, k_on_* or *k_off_*) in our model must be responsive to Wg signaling. We systematically varied one of the parameters while keeping the others constant. In all cases, varying one parameter did produce a spectrum of Sens expression levels.

We next calculated the Fano profile for each case. Only a free variation in *k_on_* produced a Fano profile that resembled the experimental data, with a Fano peak at the lowest Sens levels which dramatically declines as Sens levels increase. Thus, to recreate a Sens gradient *in silico* we kept *S_m_, D_m_, S_p_, D_p_* and *k_off_* constant and varied *k_on_* from 0.025 to 5 min^−1^. Since 1/*k_on_* defines the average time the promoter is inactive, this varied from 12 seconds to 40 minutes in our model.

#### Impact of transcription burst kinetics

Given that average time the promoter is ‘off’ is 1/*k_on_* and average time it is ‘on’ is 1/*k_off_*, we define transcription burst size and burst frequency as follows

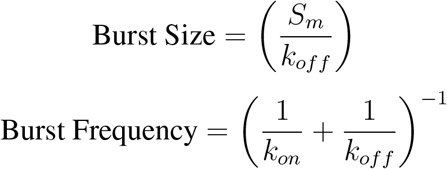

It is worth noting these values define the average burst size or frequency across exponentially distributed values. We independently varied burst size with *S_m_* (Fig. 2c) and burst frequency with *k_on_* (Fig. 2d).

As described previously, a gradient in *k_on_* can re-create the experimentally observed noise profile. Together, these observations suggest that perhaps the morphogen gradient translates into a gradient of *sens* promoter burst frequencies - at low morphogen concentrations, burst frequency is low and at high concentration, the promoter switches states rapidly. In general, we found that as promoter state switching time-scales get smaller with respect to mRNA or protein lifetimes, bursting dynamics negligibly contribute to expression stochasticity (Extended Data Fig. 5e). This is expected since frequent individual transcription bursts get time-averaged on the scale of long lived mRNA or proteins^18^.

From above, it is clear that either of *k_on_* or *k_off_* could be rate-limiting to determine burst frequency. Therefore, we also tested the effect of only varying *k_off_* while keeping the other 5 parameters constant (*k_on_* = 1/min i.e. non-limiting). Interestingly, a gradient of *k_off_* produced a very distinct Fano profile which peaked at approximately half-maximal protein expression (Extended Data Fig. 5f). *k_off_* is a coupled parameter that simultaneously affects both transcription burst size and frequency. From this we speculate that perhaps developmental genes are preferentially regulated by modulating *k_on_* rather than *k_off_* to ensure invariant protein production at higher expression levels.

After recreating the graded expression of Sens, we next sought to understand which burst parameter(s) could explain the effect of genomic position on Fano factor. Modulating burst frequency simply regenerated the noise profile seen before, as expected. Increasing burst size with *S_m_* (or even with the coupled parameter *k_off_*) mimicked the higher and larger Fano peak change as seen for *sens* at 57F5 (Fig. 4c). Thus, from simulation results, we inferred that transcription burst size for *sens* is greater at position 57F5 than at 22A3. To simulate cells of type 22A3/57F5, we simply simulated cells with two *sens* alleles with different *S_m_* values corresponding to a burst size of either 5 or 10 mRNAs. As before, alleles were simulated independent of each other to generate the Fano factor profile (Extended Data Fig. 5g).

#### Relationship between protein level and ‘constant’ Fano factor

If *k_on_* and *k_off_* are not limiting i.e. promoter switching events do not contribute significantly to expression noise; and the promoter is at 100% occupancy, the steady state protein level is described as:

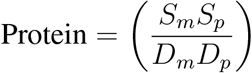

Thus, once the promoter is fully occupied, protein expression must be increased by regulating the birth-death rate constants. Correspondingly, the Fano factor will be:

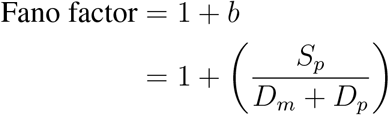

If *b* ≫ 1, then we have:

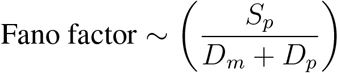

Thus the Fano factor must rise with protein level if these rate constants are perturbed. When we freely vary *S_p_, D_m_* or *D_p_* in simulations, we recreate this linear relationship (Extended Data Fig. 5d) such that if the rate constant is biased towards greater Sens protein accumulation, the corresponding Fano factor increases. We also observe signatures of a slowly rising Fano factor in our data (Fig. 3c, 4b) in the regime we describe as ‘constant’ Fano noise. We therefore speculate that *S_p_, D_m_* or *D_p_* might vary across the developmental field to expand the range of steady state Sens accumulation independent of the *sens* promoter.

### 1D fate selection model

The mathematical model for self-organization of *Drosophila* proneural tissue and fate selection is as described by Corson *et. al*.^7^ shown below

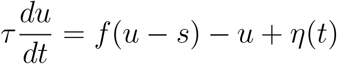

Here *u* describes the state of each cell and can range from 0 (low *u* or E-fate) to 1 (high *u* or S-fate). For our purposes, we assume u represents the fractional concentration of fate determinant Sens molecules. Inhibitory signals received from neighbor cells are summed and represented as *s* and *τ* is the time-scale of cell dynamics. All functional forms and parameter values were kept identical to Corson *et. al*.^7^ with the exception of the Brownian noise term *η*(*t*) and simplification of the model to a 1D array of competing cells with periodic boundary conditions. Pre-pattern noise was set to zero. Different levels of Sens stochasticity were simulated by running the model with the Brownian noise values drawn from distributions centred at zero, but with different standard deviations. The standard deviation in units of *u* for low, intermediate and high noise were set to 10^−6^, 10^−3^ and 10^−2^ respectively.

**Supplementary Table 1:** Sample sizes for experiments

| Figure | Sample | Sample size |
| --- | --- | --- |
| 1,2 | single tag<br>tandem tag | 19,549 cells<br>25,045 cells |
| 3 | *sens* with miR-9a sites intact<br>*sens<sup>m1m2</sup>* with miR-9a sites deleted | 50,778 cells<br>79,385 cells |
| 4 | 22A3 / 22A3<br>57F5 / 57F5<br>22A3 / 57F5 | 19,549 cells<br>27,221 cells<br>13,776 cells |
| 5 | parent stock - 22A3/22A3<br>parent stock - 22A3/57F5<br>parent stock - 57F5/57F5<br>*sens* (+ miR-9a) - 22A3/22A3<br>*sens* (+ miR-9a) - 22A3/57F5<br>*sens* (+ miR-9a) - 57F5/57F5<br>*sens* (- miR-9a) - 22A3/22A3<br>*sens* (- miR-9a) - 22A3/57F5<br>*sens* (- miR-9a) - 57F5/57F5 | **Mispatterned / Total**<br>0 / 60 wings<br>0 / 60 wings<br>0 / 88 wings<br>2 / 62 wings<br>1 / 60 wings<br>23 / 79 wings<br>3 / 83 wings<br>4 / 64 wings<br>12 / 65 wings |
| Ext. 6a | *sfGFP-sens* / *mCherry-sens*<br>*sfGFP-sens* / *mCherry-sens<sup>m1m2</sup>*<br>*sfGFP-sens<sup>m1m2</sup>* / *mCherry-sens* | 20,362 cells<br>12,975 cells<br>19,090 cells |
| Ext. 6b | Noise assay<br>FRET assay | 42,271 cells<br>42,929 cells |
| Ext. 8 | parent stock - 22A3<br>parent stock - 57F5<br>*sens* (+ miR-9a) - 22A3<br>*sens* (+ miR-9a) - 57F5<br>*sens* (- miR-9a) - 22A3<br>*sens* (- miR-9a) - 57F5 | 174 bristle pairs<br>177 bristle pairs<br>1004 bristle pairs<br>1063 bristle pairs<br>944 bristle pairs<br>605 bristle pairs |

**Supplementary Table 2:** Reaction rate constants used in simulation model

| Event | Rate constant | Value |
| --- | --- | --- |
| mRNA synthesis | $S_m$ | 0.25 mRNA/min<br>0.1-1 mRNA/min in Fig. 2c<br>0.2-0.5 mRNA /min in Fig. 4c |
| mRNA decay | $D_m^*$ | 0.0462 mRNA/min |
| Protein synthesis | $S_p^*$ | 0.9 for *sens* mRNA<br>or 1.7 proteins/min for *sens<sup>m1m2</sup>* mRNA |
| Protein decay | $D_p^*$ | 0.002 proteins/min<br>(0.01 proteins/min for simulations) |
| Promoter activation | $k_{on}$ | 0.25 - 5 events/min |
| Promoter inactivation | $k_{off}$ | 0.05 events/min<br>0.025 - 3 events/min in Ext. Fig. 5f |

*Note:* \* indicates experimentally determined rate for wing disc *sens* expression

**Supplementary Table 3:**
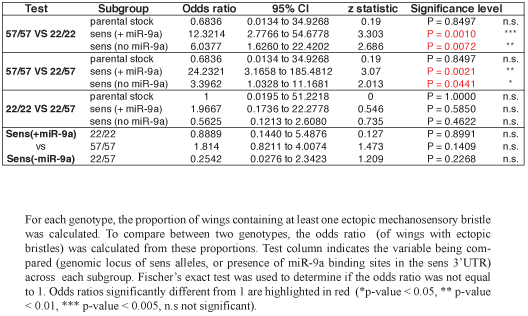
Odds ratio of wings with mispositioned mechanosensory bristles.

**Supplementary Table 4:** No annotated neurogenic genes are located in TADs containing either 22A3 or 57F5 sens landing site.

| **Locus** | **Containing TAD size** | **Genes in TAD** | **Annotated function in sensory development** |
| --- | --- | --- | --- |
| 57F5 Landing site | 11 kb | CG 10082<br>CG10321 | none<br>none |
| 22A3 Landing site | > 80 kb | CG10869<br>CG14351<br>CG31935 | none<br>none<br>none |

**Supplementary Table 5:** Sens positive cells labeled by category, assuming 1 RFU = 10 nM

| **Expected cell type** | **Single channel Sens (FCS)** | **Total Sens (FCS)** | **Expected RFU** |
| --- | --- | --- | --- |
| S-fated | Above 125 nM | Above 250 nM | Above 25 RFU |
| 1° or 2° neighbors | 25 - 125 nM | 50 - 250 nM | 5 - 25 RFU |
| Distant neighbours | Below 25 nM | Below 50 nM | Below 5 RFU |

**Supplementary Table 6:**
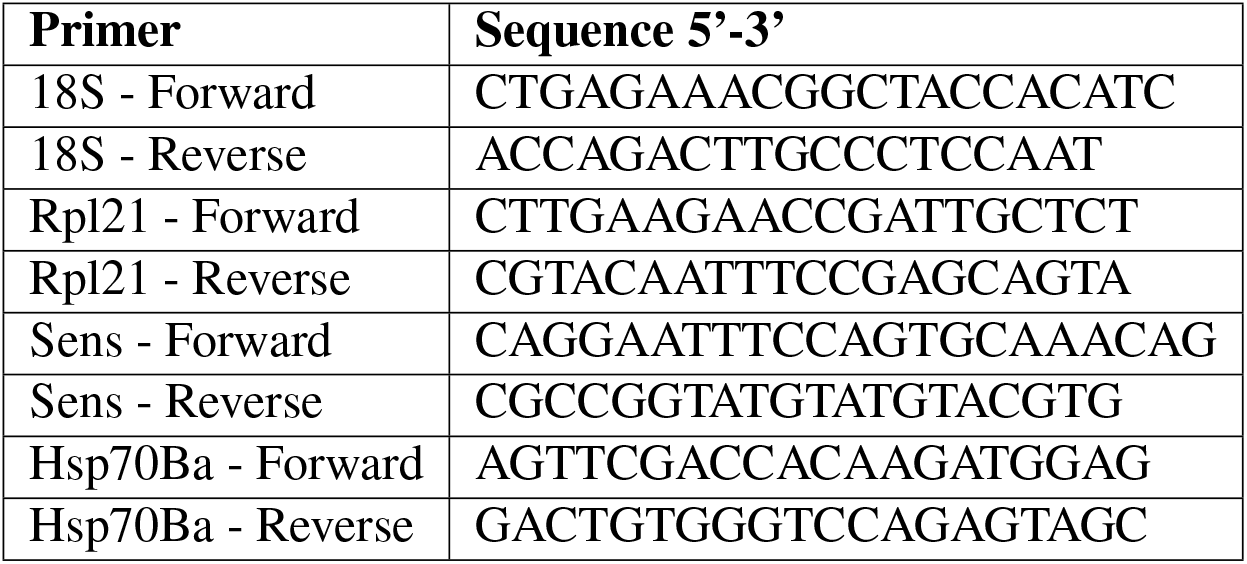
Primer sequences used for qPCR analysis

